# Conformational flexibility of tubulin dimers regulates the transitions of microtubule dynamic instability

**DOI:** 10.1101/2025.06.30.662375

**Authors:** Maddy Shred, Nicholas E. Vangos, Andrew N. Bayne, Sofía Cruz Tetlalmatzi, Wang Peng, Jean-François Trempe, David Sept, Michael A. Cianfrocco, Gary J. Brouhard

## Abstract

Microtubules are highly conserved polymers of αβ-tubulin dimers that undergo dynamic instability. While dynamic instability is conserved across eukaryotes, many of its associated conformational changes, like lattice compaction and twist, are not. Tubulin dimers sample multiple conformations in solution and undergo conformational changes during polymerization; the rate and extent of these changes describes their “conformational flexibility.” Here, we investigate the relationship between the conformational flexibility of tubulins and the dynamic phenotypes of the microtubules they produce with a comparative study of *Drosophila melanogaster* tubulin (Dm-Tb) and *Bos taurus* brain tubulin (Bt-Tb). While these tubulins share high sequence and structural similarity, their microtubules display divergent dynamic phenotypes *in vitro*, with altered transition frequencies between phases of growth and shrinkage. Dm-Tb microtubules showed drastically longer lifetimes, lower barriers to nucleation, and a high rescue frequency, while maintaining a similar growth rate to Bt-Tb microtubules. 3D reconstruction of mature Dm-Tb microtubules showed high structural conservation with mammalian microtubules. However, when we performed molecular dynamics simulations of free tubulin dimers, we found Dm-Tb to be more rigid and adopt fewer conformational states than Bt-Tb. Biochemical characterizations experimentally confirmed this finding, leading us to hypothesize that differences in the conformational flexibility of tubulins may tune the frequency of transitions between the dynamic phases of microtubules, thereby altering their stability and overall dynamic phenotypes.

## Introduction

Dynamic instability is the stochastic switching of microtubules between phases of growth and shrinkage (Mitchison and Kirschner, 1984), a process that is harnessed by eukaryotic cells to generate forces and adapt the cytoskeletal network to meet cellular demands (Desai and Mitchison, 1997; Kirschner and Mitchison, 1986). Microtubules are protein polymers composed of αβ-tubulin dimers, which join end-to-end in protofilaments that associate laterally to form a hollow, tubular lattice (Nogales, 2001). Microtubules are conserved in all eukaryotic cells since the last eukaryotic common ancestor (Margulis et al., 2006), and the tubulin gene family extends to all domains of life, with the prokaryotic homolog, FtsZ (Findeisen et al., 2014), and multiple archeal homologs (Wollweber et al., 2025; Yutin and Koonin, 2012). Gene duplication and diversification of αβ-tubulin paralogs has allowed many species to evolve specialized isotypes (Findeisen et al., 2014), form higher-order tubulin assemblies like axonemes and cortical microtubule arrays (Chaaban and Brouhard, 2017), and regulate the dynamics of their microtubules through expression of distinct isotypes (Bode et al., 2003). Nevertheless, αβ-tubulin remains under conservative evolutionary pressure, as evidenced by below-average mutation rates of orthologous αβ-tubulin coding sequences (Little et al., 1981) and the near-ubiquity of the canonical 13 protofilament (pf) polymer in all eukaryotic organisms (Chaaban and Brouhard, 2017). This evolutionary pressure has constrained the structure of αβ-tubulin dimers in order to conserve key microtubule properties like polymer formation and dynamic instability (Erickson, 2007), while allowing different species (Chaaban et al., 2018; Detrich et al., 2000; Hirst et al., 2020), and different cell types within a species (Vemu et al., 2017), to produce microtubules with distinct dynamic phenotypes.

Under the light microscope, microtubule ends appear to grow steadily before switching suddenly and stochastically to rapid shrinkage (Walker et al., 1988). At the molecular level, this simple stochastic behavior is governed by the complex mechanics of the αβ-tubulin dimer. In solution, αβ-tubulin dimers are curved (Rice et al., 2008), but show “conformational flexibility”, defined here as the ability to sample numerous conformational states, undergo internal domain rearrangements, curve, flex, and twist (reviewed in (Brouhard and Rice, 2014, 2018)). During polymer growth, these flexible αβ-tubulin dimers must collide with a growing protofilament end in the right orientation and form contacts at the longitudinal interface. The growing protofilaments vary in length, splay outwards, form clusters, twist, and bend during polymerization (McIntosh et al., 2018; Simon and Salmon, 1990). Then, the dimers must quickly form lateral bonds with neighboring dimers and undergo a curved-to-straight transition as they become buried in the microtubule lattice by the next incoming dimers. The incoming tubulin dimers have GTP in β-tubulin’s exchangeable nucleotide pocket (Weisenberg et al., 1976), which is subsequently hydrolyzed to GDP. GTP hydrolysis typically lags behind polymerization and this lag creates a stabilizing “GTP cap” that protects the polymer from “catastrophe”, or the stochastic switch to rapid shrinkage (reviewed in Howard and Hyman, 2009). Impressively, dynamic instability can be reconstituted *in vitro* without any MAPs, stabilizing agents, or external factors (Walker et al., 1988; Weisenberg et al., 1976).

The mechanics of the αβ-tubulin dimer are essential to explain the microtubule’s characteristic phases of growth and shrinkage, which stem from a mechanochemical cycle involving GTP hydrolysis and conformational changes in the polymer and individual tubulin dimers (Brouhard and Rice, 2018; VanBuren et al., 2005). Microtubule growth is entropically driven, meaning that one of the primary driving forces for polymer formation is the increase in entropy created by the release of αβ-tubulin’s hydration shell at polymerization interfaces (Erickson and Pantaloni, 1981). This entropic drive is balanced against the entropic penalty for immobilizing the flexible αβ-tubulin dimer in the microtubule lattice and conditioned by the strengths of tubulin-tubulin bonds (Brady and Sharp, 1997; Grü nberg et al., 2006). As GTP is hydrolyzed and the lattice matures, the energy landscape shifts and strain energy accumulates in the lattice (Ayoub et al., 2015; Buey et al., 2006). The release of this strain following the stochastic loss of the GTP cap occurs during the sudden catastrophe and rapid shrinkage of the microtubule (Alushin et al., 2014; Lafrance et al., 2022). Computational studies suggest that strain propagation (Igaev and Grubmü ller, 2020) and changes in dimer flexibility (Igaev and Grubmü ller, 2018) are key factors in explaining dynamic instability, but we have an incomplete understanding of how dynamic instability is regulated by the mechanics of the αβ-tubulin dimer.

Many cross-species comparative studies have used αβ-tubulins from diverse eukaryotic sources and/or mutant recombinant tubulins to advance our understanding of the mechanics of the αβ-tubulin dimer (Henkin and Reber, 2025). More specifically, cross-species comparisons have facilitated the identification of key sub-domains and conformational changes involved in dynamic instability. First, tubulin dimers have flexible loops and disordered regions that contribute to the kinetics and energetics of microtubule polymerization. In *C. elegans*, the typically flexible H1-S2 lateral contact loop in α-tubulin is pre-ordered, which was shown to activate tubulin dimers for polymerization and contribute to a faster microtubule growth rate in this species (Chaaban et al., 2018). In *Xenopus* species, increased flexibility of the β-tubulin lateral loops was predicted to stabilize the lateral contacts and lead to longer lifetimes in cold-adapted *X. laevis* microtubules compared to those from *X. tropicalis* (Troman et al., 2025).

Second, β-tubulin has a central helix (H7) that shifts significantly in the curved-to-straight transition. Two mutations in the β-tubulin H7 helix –Y222F and T238A – both lead to a reduction in the conformational differences between GTP- and GDP-tubulins (Ayukawa et al., 2021; Geyer et al., 2015; Ye et al., 2020). In *S. cerevisae*, the terminal heterodimers were less likely to be curved and GTP-lattice character was retained in GDP-tubulin, leading to a lower catastrophe frequency and slower shrinkage rate of the microtubules (Geyer et al., 2015). In *Drosophila* tubulins, the Y222F mutation disrupted a hydrogen bond the residue makes with the flexible T5 loop in GDP-tubulin, freeing the loop, increasing the nucleation potential of the tubulin dimers, and even permitting nucleation of GDP-tubulin into polymers (Ayukawa et al., 2021).

Third, tubulin dimers undergo conformational changes in response to GTP hydrolysis that are critical drivers of microtubule dynamic instability. One such conformational change is the longitudinal compaction of tubulin dimers by approximately 2 Å(Alushin et al., 2014; Hyman et al., 1995; Lafrance et al., 2022), which may weaken the bonds between tubulin dimers in the lattice. However, longitudinal compaction may not be a conserved response to GTP hydrolysis and may not explain lattice stability. In *C. elegans* (Chaaban et al., 2018), microtubules remained expanded in the GDP state, despite the fact that *C. elegans* microtubules are more dynamic than other species. Conversely, in *Xenopus* species, microtubules do undergo longitudinal compaction; however, the *Xenopus* microtubules showed high stability despite the compacted state of their mature lattice (Troman et al., 2025). In *S. cerevisiae*, compaction was not observed as a result of GTP hydrolysis but rather as a result of the binding of the EB-family protein Bim1 (Howes et al., 2017). These cross-species results suggest that compaction may not be sufficient to explain the destabilization of the mature GDP lattice.

In this work, we sought to expand our cross-species understanding of how differences in αβ-tubulin dimer mechanics can modulate dynamic instability by comparing microtubules polymerized from native *Drosophila melanogaster* tubulins (Dm-Tb) and bovine brain tubulins (Bt-Tb) *in vitro*. We chose *Drosophila* tubulin for this investigation because *Drosophila* are evolutionarily distinct from mammals, they have complex neural and cytoskeletal networks, and they have been used as a model organism in microtubule research for decades (Jakobs et al., 2022; Moriwaki and Goshima, 2016; Raff et al., 1997; Theurkauf et al., 1986; Yan and Broadie, 2007; Zhou et al., 2023). Additionally, previous reports indicate differences in the nucleation potential and catastrophe frequency of *Drosophila* microtubules (Moriwaki and Goshima, 2016; Zhou et al., 2023).

Thus, we carefully characterized the *Drosophila* microtubule dynamic phenotype *in vitro*. We find that the “dynamic phenotype” of *Drosophila* microtubules is characterized by lower barriers to nucleation, longer lifetimes, an increased rescue frequency, and a decreased shrinkage rate compared to bovine brain microtubules. To explain this dynamic phenotype, we determined a 3D-reconstruction of dynamic *Drosophila* microtubules and investigated the properties of *Drosophila* tubulins in solution with molecular dynamics simulations and biochemical assays. We find that *Drosophila* tubulins are more rigid and occupy a reduced range of conformational states, indicating that *Drosophila* tubulins have reduced conformational flexibility.

With many evolutionary constraints imposed on tubulin sequence and structure, differences in conformational flexibility could be one way for species to fine tune their microtubule networks to fit their specific needs. Changes in the conformational dynamics of tubulin dimers may therefore be an accessible avenue for species to modify the dynamic properties of their microtubules to achieve greater regulation and functional diversity, without eliminating the key microtubule properties of self-assembly and dynamic instability.

## Results

### Purified *Drosophila* and bovine tubulins show high sequence conservation between abundant isotypes

We purified native *Drosophila* tubulin (Dm-Tb) from *Drosophila melanogaster* S2 cells using TOG-affinity chromatography (Widlund et al., 2012) and bovine brain tubulin (Bt-Tb) from fresh calf brains with a cycling protocol (Gell et al., 2011). As tubulin from native sources often contains a diversity of isotypes (Roll-Mecak, 2020), we used mass spectroscopy to characterize our purified tubulin samples (Fig. S1). The Dm-Tb samples were composed primarily of α1/3 – two isotypes sharing 99.5% sequence identity – and β1 tubulin isotypes (99.95% and 85.5%, respectively; Fig. 1B). The Bt-Tb sample was more heterogeneous; the most abundant tubulin isotypes were α1D and β4B (69.7% and 52.3%, respectively; Fig. 1B and Fig. S1). Compared to each other, the sequences of the most abundant α-tubulins in our samples share 96.7% identity and 98.7% similarity, while the most abundant β-tubulins share 96% identity and 98% similarity (Fig. S1B). All isotypes identified in the two samples contain ≥85% sequence identity to each other. We visualized the location of the divergent residues on an AlphaFold3 model of the Dm-Tb α1β1 dimer (Fig. 1A). One third of the divergent residues and most of the non-conservative substitutions were located on the tubulin C-terminal tails (CTTs), as expected (Roll-Mecak, 2020). Additional non-conservative substitutions were found at the lateral contact loops and central regions of the monomers, such as the central H7 helix in β-tubulin, and the S7 strand in α-tubulin (Fig. 1A), while conservative substitutions are distributed throughout the molecule. No divergent residues were observed at the highly conserved longitudinal polymerization interfaces (Nogales, 1999). This indicates that while the most abundant Dm-Tb and Bt-Tb isotypes share high sequence conservation overall, divergent residues are localized primarily to the CTTs, lateral loops, and the globular core.

**Fig. 1.**
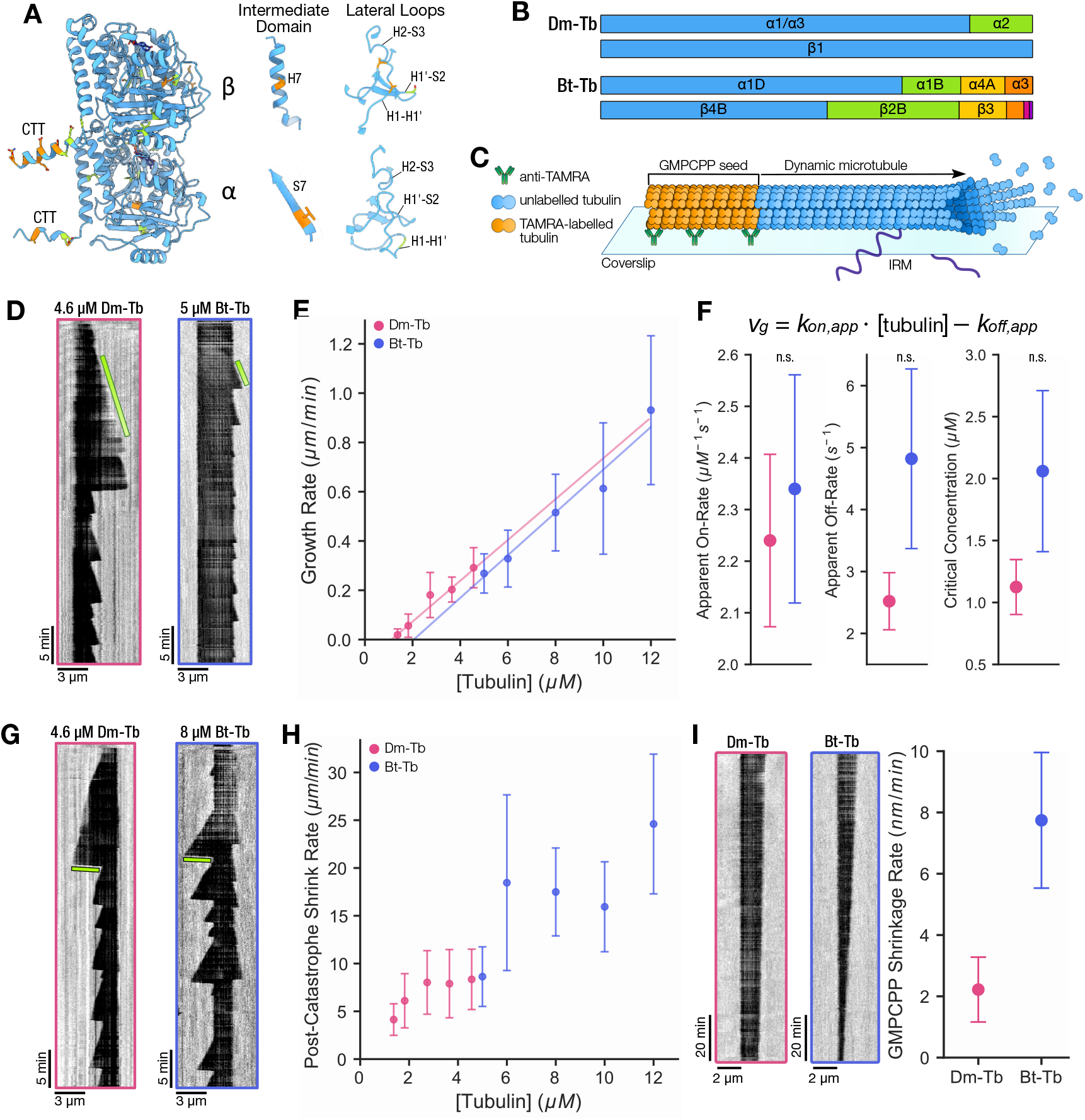
Reconstitution of *Drosophila* and bovine microtubule dynamics *in vitro*. **A)** *Drosophila* α1-β1 tubulin predicted by AlphaFold3. Residues colored by divergence of the most abundant isotypes in the study (Dm-Tb: α1-β1, and Bt-Tb: α4A-β3). Identical residues are blue, conservative amino acid substitutions are green, and non-conservative substitutions are orange. Most non-conservative substitutions are located on the CTTs (left) and other non-conservative substitutions are depicted to the right **B)** Isotype composition of the study samples, Dm-Tb and Bt-Tb, determined by mass spectroscopy (detailed in Fig. S1). **C)** Schematic of the microtubule dynamics reconstitution assay (see methods for details). **D-I)** Reconstitution of Dm-Tb and Bt-Tb microtubule dynamics at 34*°*C. **D)** Representative kymographs of microtubule growth rates for Dm-Tb and Bt-Tb. Green bars indicate the slope of the growth event. **E)** Plot of mean growth rate as a function of tubulin concentration for Dm-Tb (pink; n = 111, 123, 218, 145, 84 from ≥ 3 replicates) and Bt-Tb (blue; n = 102, 192, 196, 129, 101 from ≥3 replicates). Error bars represent the standard deviation from the mean (SD). **F)** Fitting of the data in (E) to the one-step growth curve model (above) provides the values of the apparent on-rate (*k*_*on*_), apparent off-rate (*k*_*off*_), and critical concentration (*C*_*C*_), plotted below. Error bars represent SE of the fitted parameters. Equality testing indicates that all three fitted parameters were equivalent between tubulin types (p-values ≥0.13). **G)** Representative kymographs of microtubule depolymerization rates for Dm-Tb and Bt-Tb dynamic microtubules following catastrophe. Green bars indicate the slope of the shrinkage event. **H)** Plot of mean post-catastrophe shrinkage rate as a function of tubulin concentration for Dm-Tb (pink; n = 111, 95, 175, 127, 68 from ≥3 replicates) and Bt-Tb (blue; n = 36, 116, 92, 96, 94 from ≥3 replicates). Data from the same experiments in (E) but very short shrinkage events were excluded from analysis. Error bars represent SD. **I)** Shrinkage rate of GMPCPP microtubules in the absence of free nucleotides labeled by tubulin type from ≥3 replicates each (Dm-Tb: n = 165; Bt-Tb: n = 220). Error bars represent SD.

### Reconstitution of *Drosophila* microtubule dynamics *in vitro*

To better understand the extent of the differences between *Drosophila* and mammalian microtubule dynamics, we characterized the dynamics of reconstituted Dm-Tb and Bt-Tb microtubules at 34 ^*°*^C imaged with label-free interference reflection microscopy (IRM; Mahamdeh et al., 2018). At first, we did not immediately observe a difference, as the Dm-Tb and Bt-Tb microtubules appeared to grow at similar rates. As imaging continued, however, a distinction between the two samples became clear – the Dm-Tb microtubules elongated for an extended period of time compared to the Bt-Tb microtubules. After observing these differences in the lifetimes and additional differences in the nucleation ability, we optimized the free tubulin concentrations used for Dm-Tb and Bt-Tb to allow measurement of the entire dynamic cycle, from nucleation to elongation, catastrophe, and shrinkage. Using manual kymograph analysis, we characterized the growth and shrinkage rates, lifetimes, rescue probabilities, and lag time before nucleation of Dm-Tb and Bt-Tb microtubules. The observed range of these dynamic parameters defined the “dynamic phenotype” of the microtubules. Fast nucleation, long lifetimes, and high rescue probabilities defined the Dm-Tb microtubule dynamic phenotype, which we will detail in the following sections.

Microtubule polymerization kinetics can be approximated by a linear regression model of the growth rate as a function of tubulin concentration (Erickson and O’Brien, 1992; Walker et al., 1988), allowing direct comparison of the Dm-Tb and Bt-Tb microtubules despite the different concentration ranges used. The growth curve measures the polymerization rate (*v*_*g*_) as a function of tubulin concentration, using a simple, single-pf model of the apparent on-rate (*k*_*on*_) and apparent off-rate (*k*_*off*_) of tubulin from the growing microtubule tip (Oosawa, 1975). We found that the fits produced equivalent apparent on-rate constants for Dm-Tb and Bt-Tb, at 2.24 ± 0.17 dimers·µM^*−*1^ ·s^*−*1^ for Dm-Tb and 2.35 ± 0.22 dimers·µM^*−*1^ ·s^*−*1^ for Bt-Tb (mean ± SE; p-value = 0.553; Fig. 1D-F). Despite an equivalent *k*_*on*_ to Bt-Tb microtubules, Dm-Tb microtubules show slightly increased growth rates at matched tubulin concentrations due to a shift in the critical concentration (*C*_*C*_; x-intercept of the growth curve). The *C*_*C*_ of Dm-Tb was reduced two-fold compared to Bt-Tb (1.12 ± 0.22 µM and 2.05 ± 0.65 µM, respectively; mean ± SE; Fig. 1F). Furthermore, the *k*_*off*_ for Dm-Tb was also reduced two-fold compared to Bt-Tb (2.52 ± 0.46 s^*−*1^ and 4.82 ± 1.45 s^*−*1^, respectively; mean ± SE). The reduced *k*_*off*_ for Dm-Tb indicates that while the apparent on-rate of GTP-bound tubulin dimers to the growing microtubule tip is equivalent for the two samples, GTP-bound *Drosophila* tubulins dissociate from the growing tip at a slower rate.

After measuring a two-fold reduced *k*_*off*_ for Dm-Tb, we looked for further evidence of slower dissociation of Dm-Tb dimers from the microtubule lattice. We measured the post-catastrophe shrinkage rates of Dm-Tb and Bt-Tb microtubules and found a two-fold reduced shrinkage rate overall (Fig. 1G-H). The average shrinkage rate across all tubulin concentrations was 8.6 ± 6.4 µm/min for Dm-Tb vs. 18.4 ± 9.2 µm/min for Bt-Tb (mean ± SD; n = 576 vs. 434). We observed a wide distribution of shrinkage rates, consistent with the intrinsic variability of microtubule depolymerization (Gildersleeve et al., 1992), with some variation as a function of tubulin concentration. In addition, we measured the depolymerization rate of double-cycled GMPCPP microtubules and found a three-fold slower shrinkage rate for Dm-Tb GMPCPP microtubules compared to Bt-Tb (Fig. 1I). Dm-Tb GMPCPP microtubules had a shrinkage rate of 2.22 ± 1.06 nm/min vs. 7.74 ± 2.21 nm/min for Bt-Tb GMPCPP microtubules (mean ± SD; n = 165 vs. 220). The combination of reduced shrinkage rates for both post-catastrophe and GMPCPP-stabilized microtubules with the reduced *k*_*off*_ for GTP-tubulin at the tip indicates that Dm-Tb microtubules may have stronger interactions between dimers at the tip and in the mature lattice.

### *Drosophila* microtubules display altered frequencies of transition between phases of growth and shrinkage

Next, we tested the ability of Dm-Tb and Bt-Tb to nucleate microtubules from both free subunits (i.e. spontaneous nucleation) and stabilized seeds (i.e. templated nucleation). In both cases, Dm-Tb microtubules nucleated with lower concentrations of tubulin (Fig. 2A-C). We used a pelleting assay, as previously described (Wieczorek et al., 2015), to measure the spontaneous nucleation ability and found a critical concentration of 7.35 ± 4.82 µM for Dm-Tb and 12.73 ± 2.76 µM for Bt-Tb at 36^*°*^C (Fig. 2A; Fig. S2F). In our dynamics reconstitution assay, we measured a templated nucleation threshold of 1.4 µM for Dm-Tb microtubules and 5 µM for Bt-Tb microtubules. Furthermore, the lag time before templated nucleation was reduced for Dm-Tb compared to Bt-Tb, especially at lower tubulin concentrations (Fig. 2C). Dm-Tb microtubules had a mean time to nucleate of 2.37 ± 0.05 min at 4.6 µM tubulin, compared to 11.63 ± 0.09 min at 5 µM tubulin for Bt-Tb microtubules (Exponential fit: mean ± SE). The shape of the exponential fits of the mean time to nucleate as a function of tubulin concentration indicate that the plateau of the curves were not reached in the concentration range studied; previous estimates suggest it is around 20-30 µM for bovine brain tubulin (Wieczorek et al., 2015). However, the combined nucleation data shows that Dm-Tb microtubules nucleate both faster and at lower tubulin concentrations than Bt-Tb microtubules.

**Fig. 2.**
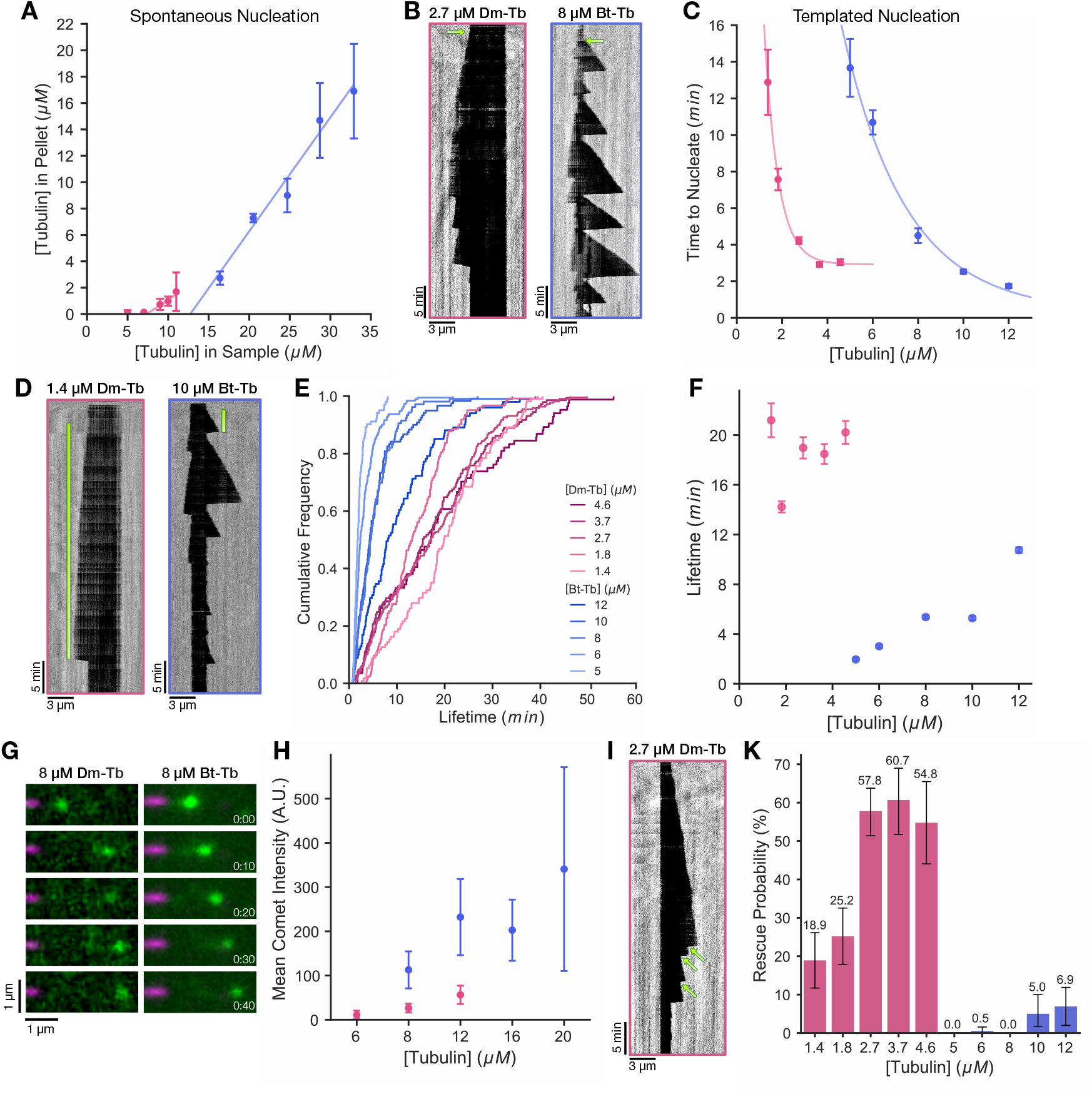
*Drosophila* microtubule dynamics show altered transition frequencies. **A)** Spontaneous nucleation of microtubules at 36*°*C. Data pooled from 3 replicates per condition. Means fit to a weighted least-squares linear regression. **B-K)** Reconstitution of Dm-Tb and Bt-Tb microtubule dynamics at 34*°*C. **B)** Representative kymographs of the nucleation lag time following addition of tubulin. Green arrows indicate the first templated nucleation event. **C)** Plot of mean time to nucleate as a function of tubulin concentration. Means calculated with an exponential fit and error bars represent the standard error of the fit. The fitted means are fit to an exponential curve: y = y0 + ae^*−kxy*^. Dm-Tb: y = 2.92 + 93.38e_*−*1.64*x*_, R^2^ = 0.998. Bt-Tb: y = 0.32 + 80.2e_*−*0.35*x*_, R^2^ = 0.990. **D)** Representative kymographs of microtubule lifetimes. Green bars represent the duration of the microtubule lifetime. **E)** Cumulative distribution function of microtubule lifetimes at each tubulin concentration. **F)** Mean microtubule lifetimes as a function of tubulin concentration. Means calculated with a gamma fit and error bars represent the standard error of the fit. **G)** Representative montage of EB3-GFP comets on Dm-Tb and Bt-Tb microtubules at 8 µM tubulin. **H)** Mean fluorescent intensity of 30 nM EB3-GFP comets on the tips of growing Dm-Tb and Bt-Tb microtubules as a function of tubulin concentration (Dm-Tb: n=123 and Bt-Tb: n=150). **I)** Representative kymograph of rescue events on Dm-Tb microtubules. Individual rescue events are indicated with a green arrow. **K)** Probability of rescue in the first microtubule growth event. No rescues were observed for Bt-Tb at 5 and 8 µM tubulin. Data presented in (B-F) and (I-K) is from the same IRM experiments presented in Fig. 1. Dm-Tb: n = 681 (84-218 per tubulin concentration). Bt-Tb: n = 621 (101-192 per tubulin concentration).

We then investigated the distribution of *Drosophila* and bovine microtubule lifetimes. The Dm-Tb microtubules had 10 × longer lifetimes than Bt-Tb microtubules at similar tubulin concentrations. We plotted cumulative distributions of the lifetimes and fit them to a Gamma distribution as previously described (Fig. 2E and Fig. S2B-D; Gardner et al., 2011; Odde et al., 1995). When polymerized at 4.6 µM Dm-Tb, the mean lifetime was 20.22 ± 0.92 min; in comparison, when polymerized at 5 µM Bt-Tb, the mean lifetime was 1.95 ± 0.09 min (mean ± SE; Fig. 2F). Even at 12 µM tubulin, the mean Bt-Tb microtubule lifetime reached only 10.74 ± 0.23 min, two times shorter than the average Dm-Tb microtubule lifetime. To investigate if the long Dm-Tb microtubule lifetimes stem from an increased GTP cap size, we reconstituted EB3-GFP comets on both Dm-Tb and Bt-Tb microtubules. The comets decorating both microtubule types were diffraction-limited spots that appeared qualitatively similar in both shape and size. However, EB3 comets on Dm-Tb microtubules showed significantly reduced mean fluorescent intensity compared to Bt-Tb microtubules across multiple tubulin concentrations (Fig. 2G-H). The dimmer EB comet intensities imply that Dm-Tb microtubules do not have extended GTP caps compared to Bt-Tb microtubules and thus that the long lifetimes must have an alternative explanation.

Next, we determined the rescue probability of Dm-Tb and Bt-Tb microtubules, and found that Dm-Tb microtubules showed significantly increased rates of rescue. Up to 60% of the Dm-Tb microtubule catastrophe events measured were followed by a rescue (Fig. 2I-K). Meanwhile, rescue events were seldom observed in Bt-Tb dynamic assays containing *<* 10 µM tubulin, and rescues occured only rarely at higher concentrations (Fig. 2K). Elevated rescue probabilities were found at all measured Dm-Tb concentrations and the rescues occured at randomly distributed locations along the entire extension of the dynamic microtubule (Fig. S2E). According to a recent study of catastrophes and rescues (Alexandrova et al., 2022), the random distribution indicates that the observed rescues are not artifacts caused by structural defects at the interface between the seed and dynamic microtubule or by surface binding effects.

Additional analysis of the microtubule minus end dynamics revealed a general conservation of the unique *Drosophila* microtubule dynamic phenotype between both the plus and minus ends. The Dm-Tb minus end had a similar growth rate, a slower shrinkage rate, a shorter lag time before nucleation, and longer lifetimes when compared to Bt-Tb minus end extensions (Fig. S3G-L). However, Dm-Tb minus ends noticeably lacked rescue events and featured fewer dynamic extensions than their plus-end counterparts, with no nucleation observed in the 1.4 and 1.8 µM tubulin conditions (Fig. S3L). Interestingly, a lack of minus end growth was also previously noted by researchers studying the nucleation of mutant and wildtype Dm-Tb microtubules (Ayukawa et al., 2021).

Finally, after measuring the microtubule dynamics of Dm-Tb and Bt-Tb microtubules at 34^*°*^C, we performed the Dm-Tb reconstitution assay at 25^*°*^C (room temperature; Fig. S3A; Goda and Hamada, 2019). *Drosophila melanogaster* are typically viable between 10-33^*°*^C (Hoffmann, 2010), meaning our original measurements were performed at the top of their viable temperature range. Relative to Dm-Tb microtubules at 34^*°*^C, we found that Dm-Tb microtubules reconstituted at room temperature showed reduced lifetimes, a longer time to nucleate, and slower shrinkage rates. However, the Dm-Tb lifetimes at room temperature remained longer than those of Bt-Tb microtubules at 34^*°*^C (Fig. S3D-F). Unexpectedly, we found that the Dm-Tb microtubule growth rates were similar at matched tubulin concentrations, producing a similar *k*_*on*_ for Dm-Tb microtubules at both 25^*°*^C and 34^*°*^C (Fig. S3C). The *k*_*on*_ of Dm-Tb at 25^*°*^C was 2.17 ± 0.057 µM^-1^s^-1^, compared to 2.24 ± 0.17 µM^-1^s^-1^ for Dm-Tb at 34^*°*^C. As Bt-Tb microtubules were previously shown to have reduced growth rates when reconstituted at lower temperatures (Chaaban et al., 2018), this finding may suggest differences in the entropic drive of polymerization that makes mammalian microtubules temperature-sensitive. Further exploration of the temperature dependence of Dm-Tb microtubules will be the subject of future investigations.

### *Drosophila* and mammalian microtubules show high structural conservation

Thus far, we measured a similar growth rate but extended lifetimes and increased rescue probabilities for *Drosophila* compared to bovine microtubules. How can the dynamic phenotypes of these two microtubule populations differ so significantly when they share *>*96% sequence identity? We searched for a structural basis of these dynamic differences with a cryo-EM study of spontaneously nucleated, undecorated, GDP-bound Dm-Tb microtubules. We determined a helically-symmetric reconstruction of a whole 14 pf *Drosophila* microtubule to a resolution of 3.5 Å(Fig. 3A). Using a novel processing pipeline, we refined the structure of a 4-dimer, 2-protofilament patch to a resolution of 2.7 Åin RELION, followed by density modification in Phenix Resolve CryoEM, resulting in a final resolution of 2.4 Å(Fig. 3B; Scheres, 2012; Terwilliger et al., 2020). Distinct lumenal loops for α-tubulin and β-tubulin demonstrate a properly sorted microtubule register in the model (Fig. 3C). The presence of electron density corresponding to GDP in the E-site confirmed that GTP hydrolysis had occurred in the dynamic Dm-Tb microtubules (Fig. 3C), and the model has a helical rise of 81.67 Å, corresponding to the compacted state of tubulin (Alushin et al., 2014; Zhang et al., 2018). Thus, this structure represents the “end-state” of the dynamic microtubule lattice following the complex processes of nucleation, elongation, and GTP hydrolysis.

**Fig. 3.**
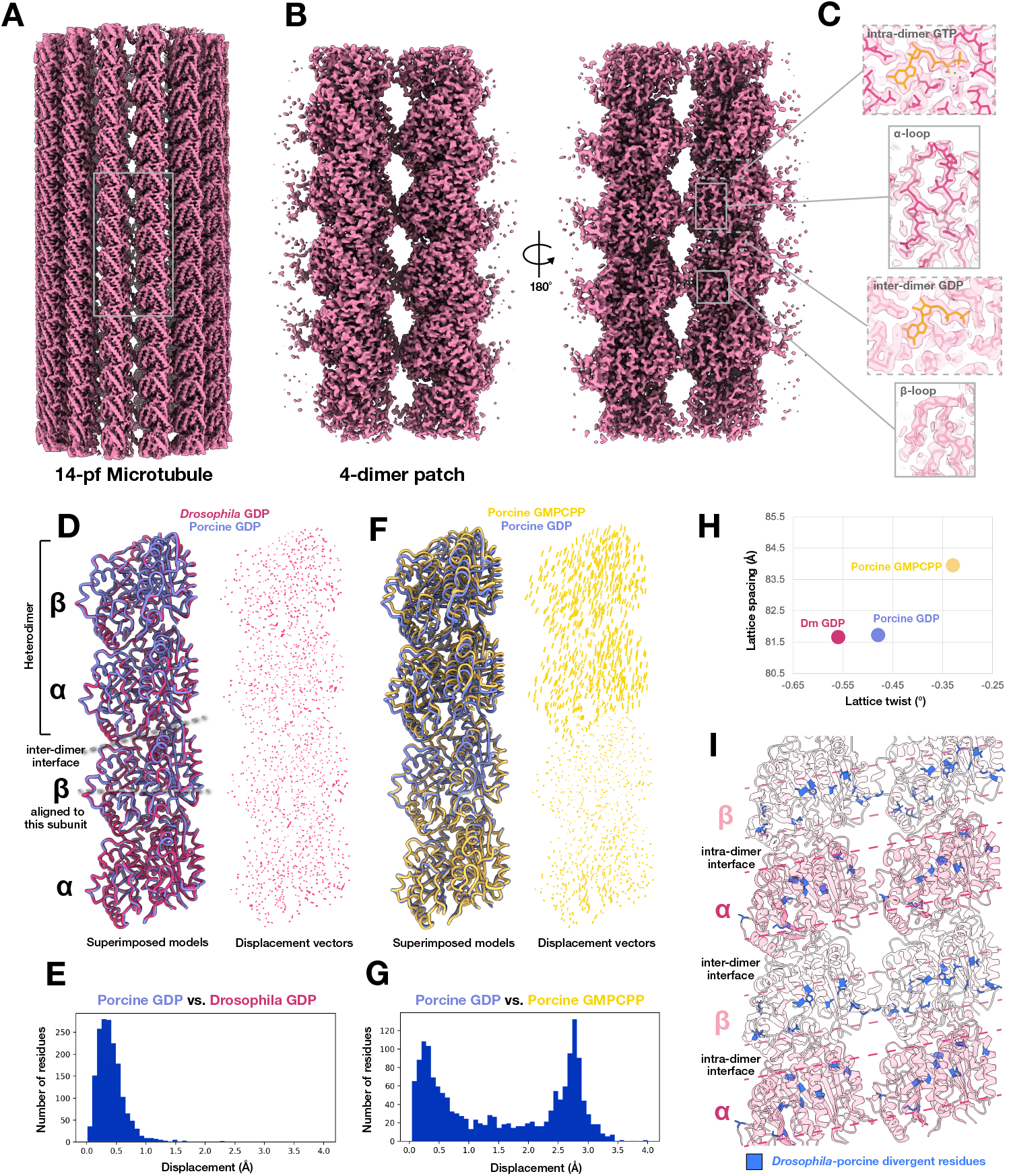
Mature 14 pf *Drosophila* and porcine microtubules share high structural conservation. **A)** Cryo-EM helical reconstruction of a mature 14 pf Dm-Tb GDP-bound microtubule. **B)** Cryo-EM single-particle reconstruction of 2×2 dimer patch of microtubule in A, front view (left) and lumenal view (right). **C)** Molecular model built into density in B clearly showing α/β tubulin register; α-tubulin is maroon, β-tubulin is pink, nucleotide is gold, magnesium is green, and coordinate water molecules are blue. **D)** Superimposed 2-dimer single protofilament models of a 14-pf GDP-bound Dm-Tb microtubule (maroon, this study) and 14-pf GDP-bound Ss-Tb brain microtubule (indigo, PDB code 6DPV: Zhang et al., 2018). Models were aligned to the central β-tubulin subunit. Residue-to-residue displacement vectors were mapped to assess conformational differences, shown to the right in maroon. **E)** Histogram of displacement vectors mapped in D. **F)** Superimposed 2-dimer single protofilament models of a 14-pf GDP-bound Ss-Tb brain microtubule (indigo, PDB code 6DPV: Zhang et al., 2018) and 14-pf GMPCPP-bound Ss-Tb brain microtubule (gold, PDB code 6DPU: Zhang et al., 2018). Models were aligned to the central β-tubulin subunit. Residue-to-residue displacement vectors were mapped to assess conformational differences, shown to the right in gold. **G)** Histogram of displacement vectors mapped in F. **H)** Plot comparing microtubule lattice parameters calculated from the 14-pf GDP-bound *Dm-Tb* microtubule (−0.56°, 81.67 Å) in maroon, 14-pf GDP-bound Ss-Tb microtubule (−0.48°, 81.73 Å) in indigo, and 14-pf GMPCPP-bound Ss-Tb microtubule (−0.33°, 83.96 Å) in gold. **I)** Ribbon model of 2×2 dimer patch from 14-pf GDP-bound Dm-Tb microtubule with residues that diverge from the Ss-Tb brain model colored cobalt blue. Dashed lines indicate central regions within each subunit that are enriched in divergent residues.

Given the significant differences we observed in the dynamic phenotypes of *Drosophila* and bovine microtubules, and that bovine and porcine microtubules are closely related, we expected to observe noticeable structural differences between mature GDP-bound *Drosophila* and porcine microtubules (Fig. 2). To our surprise, there were no noticeable structural differences in the *Drosophila* microtubule structure when compared to a previously determined GDP-bound porcine (*Sus scrofa*; Ss-Tb) microtubule structure (Zhang et al., 2018). Furthermore, both the Dm-Tb and Ss-Tb structures are directly comparable as undecorated, 14 pf, dynamic microtubules with a 3-start helix. Alignment of the two structures reveals extremely high structural similarity between the Dm-Tb and Ss-Tb microtubule structures (Fig. 3D; Zhang et al., 2018). Fewer than 3% of backbone residues shift more than 1 Åbetween the aligned models (Fig. 3E). This structural similarity becomes more apparent in contrast to the differences between the dynamic Ss-Tb structure and its counterpart GMPCPP-stabilized structure, where residue displacement vectors clearly show the ∼2 Ålongitudinal shift between compacted and expanded tubulin heterodimers (Fig. 3F-G). The dynamic Dm-Tb and Ss-Tb structures also have similar, albeit not identical, microtubule lattice twists (−0.56^*°*^ and -0.48^*°*^, respectively) with increased twist than the GMPCPP-stabilized Ss-Tb structure (−0.33^*°*^; Fig. 3H; Zhang et al., 2018).

To dissect why *Drosophila* and mammalian microtubule dynamics are so different, we revisited the localization of divergent residues in the Dm-Tb dimer (Fig. 1A). As there are minor differences in the sequences of Bt-Tb and Ss-Tb tubulins, we mapped the divergent residues between Ss-Tb and Dm-Tb tubulins onto our 4-dimer patch model from *Drosophila* (Fig. 3I). These divergent residues are enriched towards the middle of each tubulin monomer: in the globular core and lateral contact sites. However, the divergent residues are notably absent at both intra-dimer and inter-dimer longitudinal contact sites, as previously observed in the comparison to Bt-Tb tubulins and in tubulins from Antarctic fishes with highly stable microtubules (Detrich et al., 2000). While the non-conservative substitutions may alter the charge distribution, hydrogen bonding, and stability of the tubulin-tubulin interactions in the Dm-Tb microtubules, the structural similarity of the α-carbon backbones of the two models suggests they do not greatly alter the overall end-state structure of the lattice. This may suggest that more subtle, potentially allosteric changes may alter the stability of the lattice and lead to the altered dynamic phenotype of Dm-Tb microtubules.

Nevertheless, while we did not observe major structural differences in mature 14 pf, 3-start lattices of *Drosophila* and porcine microtubules, we noticed a difference in their preferred protofilament distributions. Spontaneously nucleated microtubules from different tubulin sources have been shown to adopt varying distributions of protofilament numbers (N) and helix start numbers (S) (Chaaban et al., 2018; Sui and Downing, 2010; Ti et al., 2018). Therefore, we determined the distribution of lattice geometries in our Dm-Tb microtubule sample, to learn about the preferred configuration of the lateral contacts. As microtubules with a particular protofilament number adopt different geometries when they have different helix start numbers, we will describe the lattice geometry as a combination of the protofilament number and the helix start number for specificity (e.g. N S).

We determined the distribution of microtubule geometries using layer line analysis of their Fourier transforms aided by TubuleJ (Chrétien and Fuller, 2000; Chrétien et al., 1996; Ku et al., 2020). Most Dm-Tb microtubules were composed of 14 or 15 pfs (65.0% and 24.1%, respectively; Fig. S6). 15 pf microtubules can adopt a 3-start or 4-start helix *in vitro* (Chrétien and Fuller, 2000), with differing estimates on the typical ratio (Chrétien and Wade, 1991; Sui and Downing, 2010). However, 90.9% of the 15 pf Dm-Tb microtubules adopted a 3-start helix in our sample. Both 15 3 and 14 3 microtubules feature a left-handed twist along the microtubule axis and together consist of 86.9% of the microtubules measured (Fig. 3A). In contrast, spontaneously nucleated porcine microtubules are reported to have majority 14 3 and 13 3 lattices (Zhang et al., 2018) and do not show the same enrichment of high pf number geometries. The high pf number distribution and the dominance of the left-handed twist observed in our *Drosophila* microtubule sample is similar to the reported lattice types found in highly stable *Xenopus* microtubules (Troman et al., 2025). The enrichment of left-handed 14 3 and 15 3 lattices combined with the increased left-handed twist determined in our 3D reconstruction indicates a preference for left-handed twists in *Drosophila* microtubule lattices, which may signal an altered configuration of the lateral contacts. Thus, while the end-state structures of 14 3 *Drosophila* and porcine microtubules are highly similar, the processes of nucleation and elongation lead to microtubule populations with different lattice geometry distributions. This may indicate that the dynamic differences observed between *Drosophila* and mammalian microtubules may stem from a structural or energetic difference relevant earlier in the microtubule’s dynamic cycle.

### Dm-Tb dimers explore a reduced conformational range *in silico*

The high similarity of the end-state GDP-bound structures of *Drosophila* and porcine microtubules indicated to us that the source of the unique Dm-Tb dynamic phenotype must be present earlier in the maturation process. To dig deeper into this possible source of the dynamic differences, we investigated how the the conformational states and flexibility of the tubulin dimers might differ in solution using both computational and experimental methods. First, we used all-atom molecular dynamics simulations of GDP-bound Dm-Tb and Bt-Tb dimers in solution to survey differences in their molecular motions and preferred conformational states (Fig. 4). To ensure we had good sampling and statistics, we performed multiple independent replicates of each system totaling more than 6 µs of simulations. To analyze the conformational states sampled in our simulations, we performed principal component analysis (PCA) to determine the large-scale motions of each dimer, using Bt-Tb as the reference state (Fig. 4A-B). We found that the first and second principal components (PC1 and PC2) explained 62% of the motion in Bt-Tb. The Bt-Tb dimer showed a wide range of motion and evidence of multiple distinct conformational states in PC1 and PC2. Conversely, the Dm-Tb dimer shows only a single, constrained state in the same conditions (Fig. 4A-B). PC1 and PC2 represent a mix of large-scale bending and twisting motions, as depicted in Fig. 4C-D and Supplemental Movie S1. Comparing the relative coverage of PC1 and PC2 for each dimer with the standard deviation of the PC plot, the Dm-Tb dimer explores approximately 22% of the conformational range explored by the Bt-Tb dimer on PC1 and 39% on PC2.

**Fig. 4.**
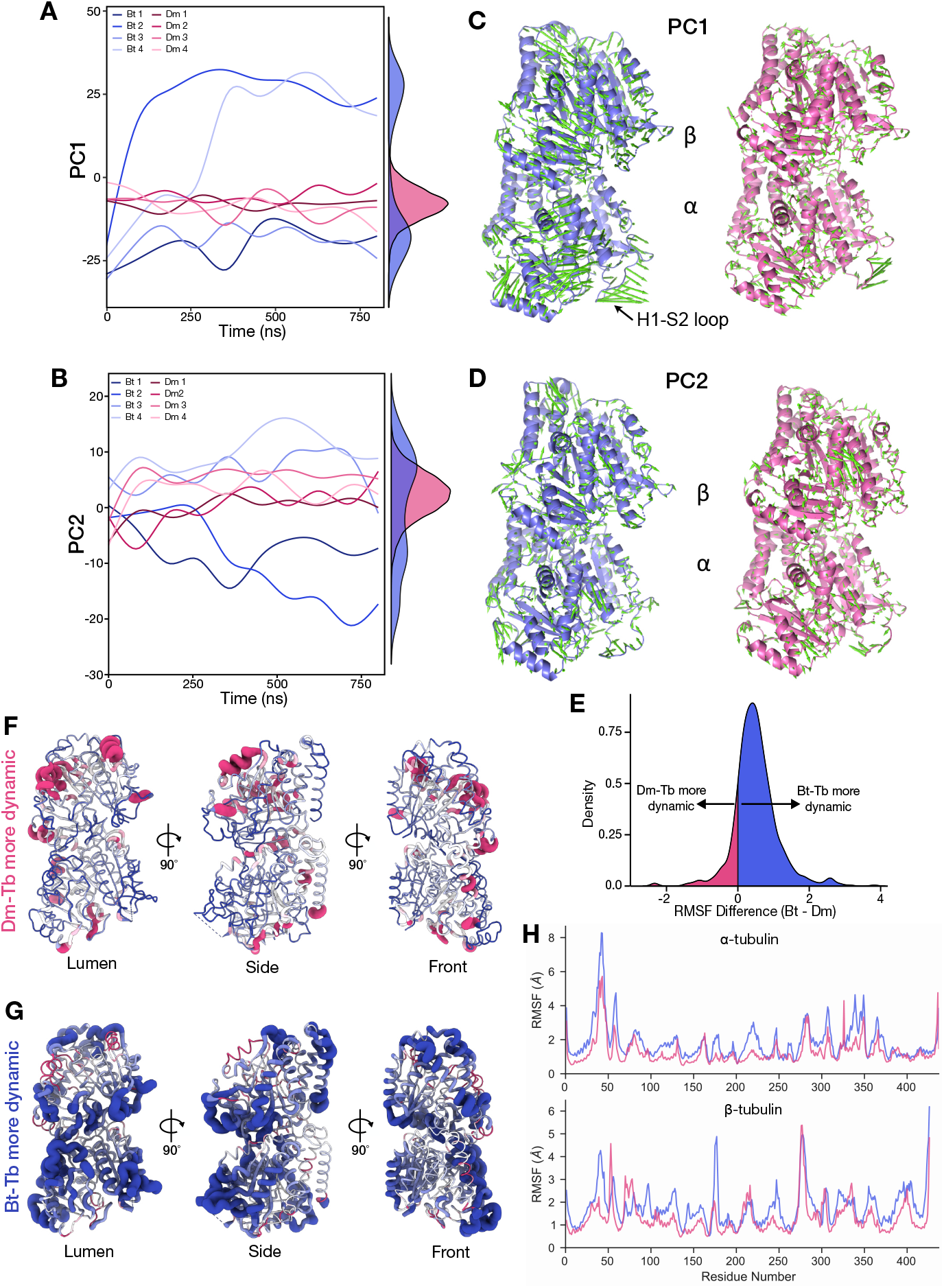
*Drosophila* tubulins display high rigidity in molecular dynamics simulations. **A-B)** Projection of the Dm-Tb and Bt-Tb all-atom MD simulations on to PC1 (A) and PC2 (B) over time. The lines show the individual simulations and the density plots on the right show the collective range of motion for each protein. **C-D)** Range of motions depicted by PC1 (C) and PC2 (D) for Bt-Tb and Dm-Tb. A model of one extremity of the specified principal component derived from the MD simulations is depicted with green vectors representing the motion across the entire principal component space. **E)** The distribution of per-residue differences in root mean-squared fluctuation (RMSF) for Dm-Tb and Bt-Tb. Positive values indicate that Bt-Tb is more dynamic and negative values indicate that Dm-Tb is more dynamic. **F-G)** RMSF differences projected onto tubulin dimer models. Coloured by RMSF difference score. The most dynamic regions in Dm-Tb (F) and Bt-Tb (G) are emphasized by increased thickness of the model. **H)** RMSF plots for α-tubulin (top) and β-tubulin (bottom) showing fluctuation of each α-carbon in Dm-Tb (pink) and Bt-Tb (blue).

To identify mobile residues and flexible regions in the tubulin dimers, we calculated the root mean-square fluctuation (RMSF) for the α-carbon of each residue in Dm-Tb and Bt-Tb (Fig. 4E-F). Since there are no insertions or deletions in the tubulin sequences, we could directly compare the dynamics of each residue by subtracting the Dm-Tb RMSF from the Bt-Tb RMSF. We found that 87% of residues were less dynamic in the Dm-Tb dimers than the Bt-Tb dimers overall (RMSF difference score *>* 0; Fig. 4G), with slightly more in α-tubulin (91%) than β-tubulin (83%). Mobile residues were widespread throughout the molecule, consistent with the highly allosteric nature of tubulin (Geyer et al., 2015; Ye et al., 2020). In α-tubulin, many of the residues with the largest RMSF difference scores were located in the H1-S2 loop (Fig. 4F). The reduced flexibility of this loop in Dm-Tb dimers compared to Bt-Tb may influence the formation or stability of lateral contacts when the dimers polymerize. In β-tubulin, the largest RMSF difference scores were located in the T5 loop, whose “in” and “out” conformations have been shown to mediate the nucleation and polymerization ability of tubulins, as GTP-binding favors the polymerization-competent “out” conformation (Ayukawa et al., 2021). The largest RMSF difference score in β-tubulin was found in residue β:D177, which forms a hydrogen bond with β:Y222 and locks the loop in the polymerization-blocking “in” conformation. A reduction in the flexibility of this loop or a difference in its preferred conformation in Dm-Tb dimers may alter the *Drosophila* microtubule’s nucleation and polymerization abilities.

Furthermore, while the Dm-Tb was less flexible overall, the dimer had a few regions that showed increased flexibility compared to Bt-Tb. In α-tubulin, the majority of the most dynamic residues in Dm-Tb were found in helix H10 near the interdimer interface and helix H12 near the CTT. Additionally, in β-tubulin, Dm-Tb was more dynamic in the region between the start of β-strand S2 to the end of α-helix H2, an area with E-site nucleotide contacts at the interdimer interface (Lö we et al., 2001). Together, these findings indicate that Dm-Tb dimers in solution may adopt a reduced conformational range overall, but show increased flexibility of certain residues on the longitudinal polymerization interfaces, which could potentially modify the Dm-Tb’s polymerization dynamics.

### Dm-Tb dimers are more stable and less flexible in solution

To dig deeper into the findings of our MD simulations, we set out to test the rigidity and occupancy of known tubulin dimer conformational states experimentally with biochemical assays. First, to measure the thermal stability of the two tubulin types in solution, we used nano-differential scanning fluorimetry (nano-DSF). We recorded the unfolding ratio (A_350_/A_330_) of the tubulin samples as the temperature was gradually increased and calculated the melting point and the onset of melting temperature, at which unfolding begins. We found that Dm-Tb dimers have a higher melting point than Bt-Tb dimers in solution (Fig. 5A-B). The melting point was 57.28 ± 0.036^*°*^C for Dm-Tb and 56.35 ± 0.033^*°*^C for Bt-Tb; a similar 1^*°*^C difference in the melting point is observed between native and Taxol-bound tubulin dimers (Morishita and Nurse, 2021), a microtubule binding drug known to increase tubulin dimer flexibility (Mitra and Sept, 2008). Interestingly, the unfolding of Dm-Tb also began at a lower temperature than Bt-Tb (Fig. 5C), suggesting there may be additional labile elements in the dimer, like the longitudinal residues showing increased flexibility in our MD simulations (Fig. 4H). The onset of melting temperature was 39.16 ± 0.20^*°*^C for Dm-Tb and 41.05 ± 0.076^*°*^C for Bt-Tb. Overall, as more energy was required to break 50% of the non-covalent bonds in Dm-Tb than Bt-Tb, they are predicted to contain more intramolecular interactions and a tighter fold in solution, congruent with the increased rigidity of Dm-Tb predicted by MD.

**Fig. 5.**
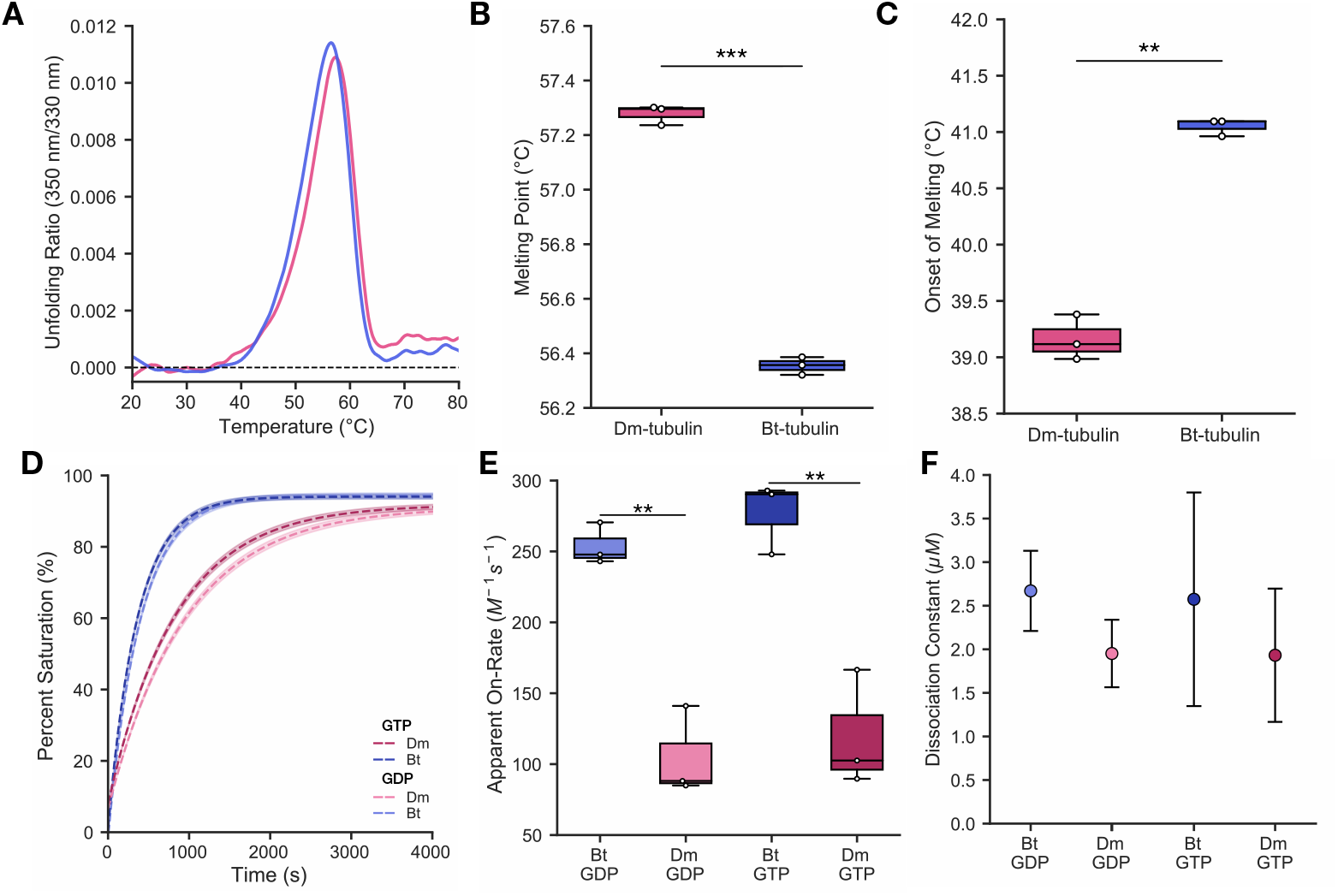
*Drosophila* tubulin dimers show increased stability and reduced conformational flexibility in solution. **A-C)** Melting point analysis by nano-differential scanning calorimetry for Dm-Tb (pink) and Bt-Tb (blue). **A)** Representative melting curves for 1 mg/mL tubulin in BRB80 buffer. **B)** Calculated melting points, the temperature at which half of the protein is unfolded (Student’s t-test: p-value = 5.6 × 10_*−*_^6^). **C)** Observed onset of melting temperature, the temperature at which unfolding of the protein began (Student’s t-test: p-value = 1.4 × 10_*−*_^3^). Melting curve data in (A-C) is from 3 measurements per tubulin type. **D-F)** Allocolchicine binding assay to measure kinetic association and affinity of allocolchicine to Dm-Tb (pink) and Bt-Tb (blue) dimers bound to either GDP (light) or GTP (dark). **D)** Kinetic association of allocolchicine to tubulin dimers in solution at 25*°*C. Data was fit to a single exponential curve (F_t_ = F_max_ - Ae^-α^_t_). The presented curve is tan average of the fits from individual kinetic measurements. The fit is represented by dashed lines and the standard error is indicated by transparent solid lines. **E)** Apparent on-rate (*k*_*on*_) from the kinetic binding curve of each individual experiment. Student’s t-test results are reported for cross-species comparisons with matched nucleotide states. GDP-bound: p-value = 0.0062; GTP-bound: p-value = 0.0084. Within-species comparisons were non-significant (p-value *>* 0.05). Data presented in **D-E** is pooled from 3 individual measurements per condition. **F)** Dissociation constant (K_D_) of allocolchicine from Dm-Tb and Bt-Tb bound to either GTP or GDP. K_D_ was measured with an equilibrium saturation curve at t = 2 hours (n=1).

To further survey the conformational flexibility of the two tubulin dimer populations in solution, we used an allocolchicine binding assay. Allocolchicine is a tubulin binding drug that emits an increased level of fluorescence when bound to tubulin (Fernholz, 1950; Fitzgerald, 1976; Hastie, 1989). The affinity of allocolchicine to free tubulin dimers has been used previously to examine differences in the conformations of tubulin in solution (Rice et al., 2008; Ti et al., 2016). Due to the location of the colchicine binding pocket at the intradimer interface of tubulin, allocolchicine is believed to only bind curved tubulin dimers, as the binding site is occluded in the straight conformation (Peng et al., 2014; Ravelli et al., 2004). As previous work showed that allocolchicine has the same affinity to both GTP- and GDP-bound tubulin dimers (Rice et al., 2008), we repeated all of our allocolchicine binding assays with both GTP and GDP, to investigate the conformational flexibility of tubulin in both nucleotide states. We hypothesized that slower conformational changes in Dm-Tb may lead to a slower capture of allocolchicine in the dimer’s colchicine binding pocket, which is only available in curved conformational states. Thus, we measured the kinetic association rate of allocolchicine to tubulin dimers, to learn about the occupancy and availability of the straight and curved conformational states (Fig. 5D-E). We found that allocolchicine bound to Dm-Tb over two times slower than it did to Bt-Tb in the same conditions. Allocolchicine bound to GTP Dm-Tb with an apparent on-rate (*k*_*on*_) of 119.59 ± 2.62 M^*−*1^s^*−*1^ while binding to GTP Bt-Tb with a *k*_*on*_ of 277.04 ± 8.937 M^*−*1^s^*−*1^ (mean fit ± SE; Fig. 5E). The apparent on-rates for binding to GDP Dm-Tb and Bt-Tb were slightly slower, at 104.71 ± 2.24 M^*−*1^s^*−*1^ and 253.78 ± 6.59 M^*−*1^s^*−*1^, respectively (Fig. 5E). The apparent on-rates of species-matched tubulins bound to GTP or GDP were not significantly different, as expected from previous studies (Rice et al., 2008). Additionally, there are no divergent residues located at the colchicine binding site of these two tubulin types and we found the dissociation constants (K_D_) for allocolchicine bound to Bt-Tb and Dm-Tb to be similar (Fig. 5F). The K_D_ was 2.57 ± 1.23 µM for GTP-bound Bt-Tb and 1.93 ± 0.76 µM for GTP-bound Dm-Tb (Fig. 5F). As previously reported (Rice et al., 2008), the K_D_ for allocolchicine bound to GDP-tubulin was equivalent to that of the same-species GTP-tubulin, at 2.67 ± 0.46 µM for GDP Bt-Tb and 1.95 ± 0.39 µM for GDP Dm-Tb (Fig. 5F). We hypothesize that the reduced *k*_*on*_ of allocolchicine to Dm-Tb is due to either reduced availability of the curved state, a slower rate of switching between curved and straight conformations, or both.

## Discussion

While dynamic instability is a conserved feature of all microtubules studied to date, microtubules from divergent species display different growth rates, lifetimes, and rescue probabilities. Our two samples, Dm-Tb and Bt-Tb, share high structural similarity and ≥96% sequence identity between the most abundant isotypes (Fig. 3, Fig. S1); yet they produce microtubules with notably distinct dynamic phenotypes. Relative to brain and human microtubules, the Dm-Tb microtubule dynamic phenotype is defined by an altered frequency of transitions between phases of growth and shrinkage: reduced barriers to nucleation, extended lifetimes, and increased rescue probabilities (Fig. 2). We propose that these differences arise from an intrinsic property of *Drosophila* tubulin dimers – reduced conformational flexibility compared to bovine tubulins (Fig. 4, Fig. 5).

Dm-Tb dimers were predicted to be more rigid and occupy a constrained conformational range compared to Bt-Tb in MD simulations (Fig. 4). While Bt-Tb dimers occupied multiple distinct conformational states, Dm-Tb dimers occupied one main conformational state in our simulations. If the dominant conformational state occupied by free Dm-Tb dimers is more similar to the lattice-bound state in terms of the twist, curvature, or internal domain arrangement, the dimers may be better suited for nucleation and elongation. Experimentally, a higher melting point indicated that free Dm-Tb dimers are more tightly folded, while slower association of allocolchicine to its binding site at the intradimer interface indicated that Dm-Tb dimers occupied curved conformational states less frequently than Bt-Tb dimers (Fig. 5). Taken together, our results indicate that Dm-Tb dimers in solution show increased rigidity, a constrained range of motion, and reduced curvature compared to Bt-Tb dimers.

Through our structural investigation, we found that 14-protofilament GDP-bound Dm-Tb microtubules were highly

Shred *et al*. | Conformational flexibility of tubulin dimers similar to 14-protofilament GDP-bound Ss-Tb microtubules. However, the samples produced different protofilament number distributions when spontaneously nucleated *in vitro*. Dm-Tb microtubules strongly favor left-handed lattice twists with 14 3 and 15 3 helices, while mammalian brain tubulins like Bt-Tb and Ss-Tb typically favor 13 3 and 14 3 helices (Chaaban et al., 2018; Zhang et al., 2018). Catalytically inactive GTP-microtubules (α:E254A mutants; Lafrance et al., 2022) and EB-decorated microtubules (Zhang et al., 2018) primarily feature left-handed lattice twists, even in “straight” 13 pf microtubules. These left-handed lattices showed reduced deformation at the seam, indicating the negative pf skew may be correlated to increased stability and reduced lattice strain (Lafrance et al., 2022). We speculate that the pf number distributions observed signal a more stable Dm-Tb microtubule population if left-handed 14 3 and 15 3 lattices are indeed more stable. Additionally, the majority of non-conservative sequence differences between Dm-Tb and Ss-Tb were located at lateral contact sites, throughout the structured core, and distal from longitudinal interfaces, implicating the divergent residues in lateral versus longitudinal contacts (Fig. 3). Differences in strain could be mediated through even slight differences in the lateral contact sites and would not necessarily be observed with our cryo-EM reconstruction. Furthermore, due to their high flexibility, the CTTs are not able to be resolved in conventional cryo-EM studies, and knowledge about the conformational states of CTTs remains elusive.

The striking similarity of our GDP-bound Dm-Tb microtubule structure to the published GDP-bound Ss-Tb structure is puzzling considering the different behaviors observed in our *in vitro* dynamics assays. Their divergent residues are notably absent at both intra-dimer and inter-dimer longitudinal contact sites (Fig. 3I). This could suggest that while the longitudinal contact sites themselves must be conserved, subtle allosteric changes throughout the tubulin body can significantly alter microtubule lattice energy and drive Dm-Tb microtubules away from catastrophe. Alternatively, these divergent residues may instead affect the conformations of soluble tubulin dimers rather than their lattice-bound conformations. This may indicate modulation of the dynamic phenotype occurs by preventing dissociation at the level of the dimer level rather than preventing catastrophe at the level of the microtubule.

In addition, the structure of the microtubule tip is known to impact dynamics but is not revealed by typical cryo-EM studies. Tubulin dimers rapidly bind to the growing ends of individual protofilaments and fall off just as rapidly if lateral bonds are not formed (Brouhard, 2015; VanBuren et al., 2005). More blunt tips have a higher average number of lateral bonds per dimer, and the average number of lateral bonds formed is predicted to be inversely proportional to tubulin’s *k*_*off*_ (VanBuren et al., 2005). Therefore, we speculate that the two-fold reduced *k*_*off*_ of Dm-Tb could potentially result in an increased average number of neighbors and reduced taper length. This would lead to quicker lateral bond formation, imparting stability to the growing lattice. Blunt microtubule ends are thought to be more stable due to less dispersal and weakening of the GTP cap (Coombes et al., 2013). Furthermore, a recent preprint combined results from cryo-ET and simulations to propose that shorter protofilament extensions and increased clustering of protofilaments favors growth by increasing the rate of favorable associations between tubulins and the growing tip (Kalutskii et al., 2025). The reduced *k*_*off*_ for Dm-Tb could result in less tapered microtubule ends that are more resistant to catastrophe, leading to the extended Dm-Tb lifetimes we observed. Specific investigation of Dm-Tb microtubule tip structures with cryo-ET would be needed to confirm or disprove this hypothesis.

Furthermore, the lateral bonds could simply be stronger in Dm-Tb lattices than Bt-Tb lattices as indicated by the post-catastrophe and GMPCPP shrinkage rates we measured (Fig. 1H-I). We wonder if the strong preference for a negative protofilament skew in Dm-Tb microtubules is indicative of stronger lateral bonds. Computational models predict that stronger lateral bonds may lead to fewer “cracks” between adjacent protofilaments, resulting in less frequent catastrophes, longer lifetimes, and a higher rescue frequency (Margolin et al., 2012), as we observe for Dm-Tb microtubules. Additionally, the lateral polymerization interfaces show an increased level of divergence across tubulin isotypes (Roll-Mecak, 2020), implicating the lateral loops in tuning microtubule dynamics. In our MD simulations, the H1’-S2 loop of *Drosophila* β-tubulin appeared to be more dynamic than in bovine tubulin (Fig. 4). In our 3D reconstruction, we saw slight differences in the positioning of this loop, as it contained the largest Ss-Tb:Dm-Tb displacement vectors, suggesting an altered conformation of the loop in mature Dm-Tb microtubules (Fig. 3D). Thus, we speculate that a more flexible H1-S2’ loop could potentially better promote or sustain lateral interactions, contributing to the long lifetimes of *Drosophila* microtubules.

While Dm-Tb microtubules are the first species reported with this specific dynamic phenotype, microtubules in other contexts that also show extended lifetimes and increased stability may hold clues to understanding the source of this dynamic divergence. For example, bovine brain microtubules reconstituted in increasing concentrations of glycerol were found to have increased lifetimes, rescue probabilities, and nucleation abilities; reduced shrinkage rates; and only a slight reduction in the growth rate (Molines et al., 2024). This was thought to be due to the micro-viscosity of glycerol reducing the rate of conformational changes in the αβ-dimer. Additionally, microtubule minus ends have longer lifetimes and an increased rescue frequency than plus ends when controlling for growth rate (Strothman et al., 2019). Strothman et al. predicted that these differences stemmed from a reduced *k*_*off*_ of tubulin at the minus end – reminiscent of the reduced *k*_*off*_ we found for Dm-Tb dimers. With such similar lattice structures between *Drosophila* and mammalian tubulins, the reduced Dm-Tb *k*_*off*_ may stem from an intrinsic property of the tubulin subunits, such as the reduced conformational flexibility we identified.

We were intrigued to find that Dm-Tb GDP microtubules were compacted, as previous studies of invertebrate microtubules nucleated *in vitro* report a lack of compaction in the absence of MAPs (Chaaban et al., 2018; Howes et al., 2017). Thus, our work showing its conservation in the invertebrate *Drosophila* indicates that compaction as an intrinsic response to GTP hydrolysis may have arisen at multiple points during the evolution of the microtubule cytoskeleton. However, destabilization as a result of lattice compaction and stability conferred by expansion does not appear to be fully conserved across species. One such invertebrate tubulin that remained expanded following GTP hydrolysis, from *C. elegans*, also produced the most intrinsically dynamic microtubules observed *in vitro* to date (Chaaban et al., 2018). These expanded GDP lattices were not particularly stable. Alternatively, our work, along with recent structural findings from highly stable *Xenopus* microtubules, shows that some compacted GDP lattices nonetheless remain highly stable (Troman et al., 2025). Therefore, it is not always true that compacted lattices are unstable while expanded lattices are stable, leading us to question whether lattice compaction is truly a molecular driving force of dynamic instability. Additionally, we hypothesize that instead of being intrinsically unstable, compaction following GTP hydrolysis is an avenue for increased regulation of microtubule stability, MAP recognition (Siahaan et al., 2022), and tubulin code modifications (Shen and Ori-McKenney, 2024; Yue et al., 2023). We speculate that this could be why it is observed in more complex organisms like vertebrates and fruit flies, but absent in organisms like yeast and *C. elegans*, which may have fewer regulatory needs.

Microtubules and their characteristic dynamic instability have evolved as a core component of eukaryotic cells since the last eukaryotic common ancestor (Margulis et al., 2006). While all eukaryotic cells harness the power of dynamic instability to perform essential functions, many organisms have fine tuned the dynamic phenotypes of their microtubule networks to suit their unique needs. Many factors can influence this dynamic phenotype, including abiotic pressures like temperature and salinity (Li and Moore, 2020), the typical cell shape and size (van Grinsven and Akhmanova, 2025), and the co-evolution of microtubules and their MAPs (Kennard et al., 2025). The different intrinsic properties of tubulins and their microtubules may therefore stem from both advantageous adaptions and epiphenomenons. However, regardless of the source of this variation, the natural diversity of the intrinsic microtubule dynamic phenotypes among divergent tubulins can be leveraged, through comparative studies like ours, to learn about tubulin as a molecular machine.

## Methods

### Tubulin purification

#### *Drosophila* tubulin

Native *Drosophila melanogaster* tubulin (Dm-Tb) was purified from S2 cell culture using TOG-affinity chromatography as described by (Widlund et al., 2012). The TOG-affinity column was prepared by coupling 20 mg of GST-TOG1/2 recombinant protein expressed in Bl21(DE3) cells to a 1 mL HiTrap NHS-activated column for 4 hours at 4^*°*^C. Remaining NHS groups were blocked with 0.5 M ethanolamine and 0.5 M NaCl for 4 hours at 4^*°*^C and the prepared TOG-affinity column was stored at -20^*°*^C in 1x Phosphate Buffered Saline (PBS; 137 mM NaCl, 2.7 mM KCl, 10 mM Na_2_HPO_4_, 1.8 mM KH_2_PO_4_; pH 7.4) with 50% glycerol. Liquid cultures of *D. melanogaster* Schneider’s S2 cells were grown in spinning flasks containing Schneider’s medium supplemented with 10% heat-inactivated fetal bovine serum and 0.5 mg/mL penicillin-streptomycin-glutamine at 25^*°*^C. Confluent cultures were centrifuged at 900xg to harvest the cells. Cell pellets were stored at -80^*°*^C until the day of purification.

Approximately 3-4 g of S2 cell pellets from approximately 1-1.3 L of cell culture were used for each purification and Dm-Tb from 8 separate purifications was used in this work. S2 cell pellets were resuspended in BRB80 buffer (80 mM PIPES, 1 mM MgCl2, 1 mM EGTA; pH 6.9) with cOmplete™, EDTA-free protease inhibitors and lysed with a Dounce tissue grinder on ice. The lysate was clarified by ultracentrifugation at 80,000 rpm for 15 minutes and passed through a 0.22 µm syringe filter. Clarified lysate was loaded onto the TOG-affinity column equilibrated in BRB80 at 0.5 mL/min. Depending on the components of the chromatography system used, the lysate was either circulated continuously over the column for one hour with a sample pump or manually re-applied an additional 2-3 times using an injection loop, to maximize Dm-Tb binding to the TOG domains. Three BRB80 wash steps were used: (1) 20 column volumes (CV) BRB80 with 100 µM GTP at 0.5 mL/min, (2) 40 CV BRB80 with 10 µM GTP at 1 mL/min, and (3) 20 CV BRB80 with 0.1% Tween20 and 10% glycerol at 1 mL/min. The bound Dm-Tb was eluted with 500 mM ammonium sulfate in BRB80 buffer with 10 µM GTP at 1 mL/min. The eluted protein was immediately desalted into BRB80 storage buffer using a PD10 desalting column. The BRB80 storage buffer contained different supplements depending on the intended downstream use of the Dm-Tb: 10 µM GTP and 5% glycerol for microscopy experiments, 10 µM GTP for cryo-EM experiments, or no supplements for allocolchicine binding assays. In one preparation, an additional purification step using a cation exchange column (HiTrap SP FF 1 mL) was used before desalting the tubulin into storage buffer. No improvement in purity was noticed with the cation exchange column, while a reduction in the flow rates of the wash steps was seen to reduce the purity of the final Dm-Tb samples. The purified Dm-Tb was concentrated to working concentrations using a centrifugal filter depending on estimated yeild (Amicon-15 10K for a final volume of 250 µL or Amicon-4 30K for a smaller final volume), flash frozen in small aliquots with LN_2_, and stored at -80^*°*^C. The TOG-affinity column was washed with 10 CV 5x PBS at 1 mL/min and 10 CV 1x PBS with 50% glycerol at 0.2 mL/min before being stored at -20^*°*^C.

#### Bovine brain tubulin

Bovine brain tubulin (Bt-Tb) was purified from fresh calf brains through two cycles of polymerization and depolymerization as described previously (Ashford and Hyman, 2006; Gell et al., 2011). Briefly, the fresh brains were homogenized with a blender at 4^*°*^C and clarified by centrifugation and filtration with cheese cloth. Two purification cycles were performed with each of the following steps: (1) microtubule polymerization at 35^*°*^C, (2) isolation of the polymer by centrifugation at 35^*°*^C, (2) microtubule depolymerization at 4^*°*^C by douncing, and (4) removal of debris by centrifugation at 4^*°*^C. After the second cold depolymerization, the tubulin was further purified on a column containing Fractogel EMD SO_3-_ (M) and the eluted Bt-Tb was stored at -80^*°*^C. An additional cycle of polymerization and depolymerization as described above was used to isolate polymerization competent Bt-Tb from the purified Bt-Tb sample (Hyman et al., 1991). Concentrated aliquots of cycled Bt-tub was flash frozen in LN_2_ and stored at -80^*°*^C in BRB80 buffer for use in future experiments.

#### Sample purity assessment

The purity of all tubulin preparations was determined by SDS-PAGE and selected preparations were analyzed by mass spectrometry as described below. To assess the percentage purity of tubulin samples by SDS-PAGE, tubulin samples were mixed 1:1 with 2X Laemmli sample buffer and heated at 75^*°*^C for 10 minutes. 6 µg of each sample were resolved on a 4-20% Tris-Glycine gel at 120 V and were visualized using Coomassie stain. The concentration of purified tubulins was determined using a concentration curve of absorbance at 280 nm on a DS-11 FX spectrophotometer (DeNovix Inc.) and an extinction coefficient of 115,000 M^*−*1^cm^*−*1^.

### Mass spectrometry analysis of tubulin samples

For mass spectrometry analysis, tubulin samples were mixed with 6M urea / 50 mM Triethylammonium bicarbonate (TEAB; pH 8.0), reduced with 5 mM dithiothreitol (DTT) for 30 minutes at 37^*°*^C, and alkylated with 15 mM iodoacetamide at room temperature for 30 minutes in the dark. Samples were diluted to 1 M urea with 50 mM TEAB, Trypsin/Lys-C mix (Promega) was added in a 1:25 (w/w) ratio, and the digestion was incubated at 37^*°*^C for 16 hours. Trifluoroacetic acid was added to a final concentration of 0.5% to stop the digestion and samples were dried in a LabConco Centrivap DNA vacuum concentrator. Dried pellets were re-suspended in 0.5% formic acid / 5% acetonitrile and were cleaned using C18 spin columns (Pierce), according to manufacturer’s instructions. Eluates were again dried in a LabConco Centrivap DNA vacuum concentrator. Purified peptides (2 µg) were resuspended in 0.1% formic acid / 5% acetonitrile, injected onto an ACQUITY UPLC BEH C18 column (130 Å, 1.7 µm, 1 mm x 150 mm), eluted with a 90 min 5-40% gradient of acetonitrile in 0.1% formic acid at 40 µl min^*−*1^, and analysed with an Impact II Q-TOF spectrometer. Data was acquired using data-dependent auto-MS/MS with an m/z range of 150-2200, and a fixed cycle time of 3 s with active exclusion.

Raw data were processed using Andromeda, integrated into MaxQuant (version 2.1.4.0). Searches for tryptic peptides (before K,R) were performed against the *D. melanogaster* (UP000000803) and *B. taurus* (UP000009136) proteomes with a maximum of two missed cleavages and a minimum peptide length of 7 amino acids. All searches were performed with carbamidomethylation (C) as a fixed modification. Variable modifications included oxidation (M), acetylation (protein N-term, K), deamidation (NQ), and phosphorylation (STY). Default MaxQuant instrument parameters for Bruker Q-TOF data were used, including a first search peptide mass tolerance of 20 ppm, main search peptide tolerance of 10 ppm, and isotope match tolerance of 0.005 Da. The false discovery rate threshold was set to 1%. Unique and razor peptides were used for quantification. For purity estimations, iBAQ values for all tubulin protein groups were summed, divided by the total iBAQ of all identified proteins within a given run, and were expressed as a percentage. Proteins which were “identified by site”, “reverse” (decoy proteins), and “potential contaminants” were excluded from this calculation.

### Microtubule dynamics reconstitution assay

#### Experimental set up

Dynamic Bt-Tb and Dm-Tb microtubules were reconstituted in custom built flow cells containing GMPCPP-stabilized microtubule seeds as described previously (Gell et al., 2010). The GMPCPP-stabilized microtubule seeds used in all dynamic reconstitutions were prepared with two cycles of polymerization and depolymerization of 25% tetramethylrhodamine (TAMRA, ThermoFisher Scientific) labeled Bt-Tb and 75% unlabeled Bt-Tb with GMPCPP (Gell et al., 2010; Hyman et al., 1991). GMPCPP-stabilized Dm-Tb microtubules used to measure the GMPCPP-microtubule shrinkage rate were prepared in the same manner except that unlabeled Dm-Tb was used exclusively.

Glass coverslips were cleaned and silanized as previously described (Gell et al., 2010) with the exception that, after sonication in 0.5M KOH, the coverslips were treated in a plasma oven (Plasm Etch) for 3 minutes instead of treatment with piranha solution. Coverslips were stored under vacuum in a desiccator until flow cell construction. Flow cells were constructed by placing thin strips of double-sided tape between two coverslips (22 × 22 mm and 18 × 18 mm) on a custom-built housing. Anti-TAMRA antibody was attached to the surface of the flow cell with 1-2 minutes of incubation and remaining surfaces were blocked with 1% Pluronic F-127 for 20 minutes minimum. The channel was rinsed with warm BRB80 three times before flowing in the TAMRA-labeled GMPCPP-stabilized seeds. Unbound seeds were washed away with BRB80 before the channel was placed on the microscope objective.

On each day of experiments, new aliquots of tubulin were thawed on ice, sub-aliquoted into appropriate volumes for each channel, and stored in LN_2_ until the time of the experiment. Microtubules were nucleated from the GMPCPP seeds by flowing in the reaction mix, which was tubulin and reaction buffer (BRB80; pH 6.9, 1 mM GTP, 100 mM bovine serum albumin, 10 mM DTT, 250 nM glucose oxidase, 64 nM catalase, 40 mM D-glucose, and 0.05% methylcellulose). The reaction mix was prepared with the appropriate concentration of tubulin on ice immediately prior to imaging.

#### Interference reflection microscopy

The dynamics of unlabeled microtubules were imaged using interference reflection microscopy as previously described (Mahamdeh et al., 2018). Imaging was performed on a Zeiss Axiovert Z1 inverted microscope with a single 100X /1.46 NA objective (Zeiss, 420792-9800-000) and 50/50 mirror (Chroma, Bellows Falls, VT, USA). The temperature in the channel was set to 34^*°*^C with a CU-501 Chamlide lens warmer (Live Cell Instrument) unless otherwise specified. Channels performed at 25^*°*^C were conducted at room temperature with the lens warmer mounted on the objective lens with the warming function turned off to monitor the temperature and a thermometer measuring the room temperature placed next to the microscope. A LED lamp (pE-300white, CoolLED, UK) was used to illuminate the FOV with 440 nm light and the NA of the illumination was set by two aperture irises adjusted to improve the signal to noise ratio. Images were recorded on an Andor Neo sCMOS camera with a 63 nm pixel size and image acquisition was controlled by MetaMorph (Molecular Devices).

Time lapse sequences of dynamic microtubules were collected for 50-60 minutes with the use of a Definite Focus (Zeiss) to maintain image focus and a custom acquisition loop in MetaMorph. The loop acquires 40 images with a 10 ms exposure time sequentially, averages them, and saves them as a single Tiff file. The reaction mix containing tubulin was added to the channel following the start of image collection, the frame of addition was noted from the acquisition loop number on MetaMorph or by turning off the LED lamp at the moment of addition, and the channel was capped with nail polish. Time lapse images were initially collected at 5 second intervals for the 34^*°*^C conditions and then at 2 second intervals for the 25^*°*^C condition. Time lapse sequences of GMPCPP microtubules shrinking were collected at 30 second intervals over the course of 3 hours (Bt-Tb) and 4 hours (Dm-Tb). A custom Python script (brouhardlab GitHub) was used after collection to determine the precise frame rate in seconds per frame (spf) from the image file metadata. After each time lapse sequence, 200 frames were collected while moving the FOV at high speed and the median was taken to create a background image used in processing.

#### IRM dynamics movie processing and analysis

Image processing and kymograph generation were conducted in Fiji (Schindelin et al., 2012), while analysis was conducted with Python. Time lapse sequences were first flat-field corrected as described previously (Cleary et al., 2022) using a custom macro in Fiji (brouhardlab GitHub), with the exception that the processed movie was not inverted and microtubules remained represented by dark pixels. Next, the movies were corrected for stage drift with a custom drift correction macro in Fiji (brouhardlab GitHub). Final processed movies were converted to 8-bit images and cropped to remove sections of the FOV not visible throughout the entire duration of the movie due to drift correction.

Processed movies were analyzed manually using kymographs. Lines were drawn along the longitudinal axis of either all microtubules within the FOV if fewer than 100 microtubules were visible or approximately 80-100 microtubules in densely packed FOVs with over 100 microtubules. Kymographs were generated from these lines and each kymograph with a microtubule that completed at least one growth cycle in focus within the FOV was kept for analysis. A segmented line region of interest (ROI) was drawn on each kymograph from the first nucleation event until the first catastrophe event and saved for analysis. Only the first growth event from each microtubule was measured in order to analyze the time to nucleate and ensure the “stabilized seed” template contained the same blunt end structure. For each movie, an additional ROI was drawn to indicate the frame in which tubulin was added to the channel.

The growth rate, time to nucleate, growth lifetime, growth length, shrinkage rate, shrinkage lifetime, shrinkage length, and presence or absence of a rescue event were calculated using the information contained in the ROIs with Pandas and Numpy packages in a custom Python script (brouharlab GitHub). The resulting dynamic parameters were fit to Gaussian distributions (growth and shrinkage rates), exponential functions (time to nucleate), and Gamma distributions (lifetimes) with the SciPy package in Python. The fitted mean growth rates were then fit to a linear regression model:

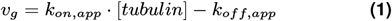

to determine the apparent on-rate (*k*_*on*_), apparent off-rate (*k*_*off*_), and the critical concentration (*C*_*C*_; *k*_*off*_*/k*_*on*_; Oosawa, 1975) using weighted least-squares regression in Python (statsmodels). The SE of the *C*_*C*_ was calculated using error propagation from the SE of the WLS fits of the *k*_*on*_ and *k*_*off*_. The fitted time to nucleate means were fit to an exponential curve using the SciPy package in pythonfor the growth rate and to an exponential curve for the time to nucleate data were then conducted with Python module statsmodels and the SciPy package, respectively.

### Pelleting assay for spontaneous nucleation

To measure the critical concentration for spontaneous nucleation of Dm-Tb and Bt-Tb, we performed a pelleting assay adapted from previous studies (Mitchison and Kirschner, 1984; Wieczorek et al., 2015). Tubulin at concentrations ranging from 5-11 µM for Dm-Tb and 16.4-32.9 µM for Bt-Tb was incubated on ice for 5 minutes in BRB80 with 1 mM GTP. The samples were placed in a heating block and incubated at 35^*°*^C for 1 hour to allow microtubules to polymerize. The solution was centrifuged to pellet microtubule polymer at room temperature for 5 minutes in a Beckman airfuge at 30 psi. The supernatant was removed from the centrifuge tube and the pellet was washed once with warm BRB80. The pellets were incubated on ice for 10 minutes and resuspended in cold BRB80. Dm-Tb samples were directly mixed 1:1 with 2× SDS-PAGE loading buffer while the Bt-Tb samples were diluted 1:1 with BRB80 before being mixed 1:1 with 2× SDS-PAGE loading buffer. The samples were boiled for 8 minutes at 90^*°*^C and loaded onto a 4-12% SDS-PAGE gel (GenScript). Four tubulin samples of known concentration were included on each gel as loading controls for future analysis. The gel was run at 140 V for 1 hour and 5 minutes, rinsed with water, and stained with Coomassie for 2 hours, and destained in a solution of 10% acetic acid overnight for 16-18 hours. The gel was scanned and imported into Fiji for analysis. The intensity of the bands was analyzed with the “Gels” analysis tool in Fiji and the concentration of tubulin in the pellet was determined by comparison to the tubulin loading controls. The x-axis intercept was calculated from a line of best fit to all points *>* 0.1 µM to determine the critical concentration for spontaneous nucleation at 35^*°*^C.

### Cryo-EM of undecorated *Drosophila* GDP-bound microtubules

#### Sample preparation

Undecorated, dynamic Dm-Tb microtubules were prepared by spontaneous nucleation and polymerization of purified Dm-Tb at 36^*°*^C in the presence of GTP. 10 µM Dm-Tb was incubated in BRB80 supplemented with 1 mM GTP and 1 mM MgCl_2_ for 10 minutes in a thermocycler. Due to the presence of only few, extremely long microtubules, the microtubules were sheared by pipetting when removed from the thermocycler and applied to the grids within 1-5 minutes. Observation by IRM confirmed that the microtubules remained stable and increased in number following shearing. A new preparation of microtubules was polymerized for each group of 2-3 grids prepared.

C-flat™ holey carbon-coated grids (R2/1, Electron Microscopy Sciences) were glow discharged for 15 seconds at 10 mA, no more than 15 minutes before use (Pelco easiGlow system). Prepared microtubules were loaded onto each grid in 4 µL volumes and incubated for 1 minute. The grids were blotted and plunge-frozen in liquid ethane with an FEI Vitrobot MkIV Grid Plunging System. The humidity of the Vitrobot was set to 100% and the temperature was 25^*°*^C. Prepared grids were transferred to storage pucks and kept in LN_2_ until imaging. Prepared grids were imaged at two facilities. Approximately 60 single-frame micrographs were collected solely for layer line analysis during grid screening at the first facility. The majority of the data, in the form of super-resolution videos, was subsequently collected from the quality-controlled grids for 3D reconstruction; the resulting micrographs were used for both 3D reconstruction and layer line analysis.

#### Image collection for TubuleJ Analysis

Micrographs used solely for layer line analysis were imaged at the Facility for Electron Microscope Research at McGill, on a 200 kV Cryo-STEM microscope (FEI Tecnai G2 F20) with a cryo-holder from Gatan. Micrographs were collected of intact microtubules observed at a nominal magnification of ×62,000 with a defocus range of −2 µm to −3.5 µm. Single-frame images were acquired with a 16 MP CMOS camera (TVIPS TemCam XF416) with a pixel size of 1.76 Å. Micrographs with no ice contamination and clear, uninterrupted microtubules with sections of 200 nm or greater were selected for analysis.

#### Image collection for 3D Reconstruction

A total of 5,104 super-resolution videos were collected at the University of Michigan Cryo-EM Facility at a nominal magnification of ×105,000 on an FEI Titan Krios G4i microscope (Thermo Fisher), equipped with a K3 direct electron detector and BioQuantum energy filter (Gatan) and set to a slit width of 20 eV. Collection was performed semiautomatically using SerialEM (Mastronarde, 2005) at a dose rate of 13.0 e^*−*^ pixel^*−*1^ s^*−*1^ for a total dose of 50 e^*−*^/Å_2_ over 50 frames with a defocus range of −0.5 µm to −2.0 µm. The electron microscopes used undergo regular and frequent re-calibration by solving the structure of apoferritin and adjusting the pixel size accordingly (as in Montecinos et al., 2025).

#### Image processing and 3D-reconstruction

All following steps were performed in RELION-5.0-beta (Kimanius et al., 2021, 2016; Scheres, 2012; Zivanov et al., 2018) with all 3D steps using blush regularisation unless otherwise noted (Kimanius et al., 2024). Dose-fractionated image stacks were motion-corrected, dose-weighted, and 2*×* binned to the physical pixel size of 0.832 Åby MotionCor2 (Zheng et al., 2017). The contrast transfer function (CTF) of dose-weighted, motion-corrected micrographs was estimated by CTFFIND-4.1.14 (Rohou and Grigorieff, 2015). Micrographs with CTF fit resolution poorer than 6.5 Åand excessive ice contamination were removed, resulting in a final total of 4,448 micrographs. Selected micrographs were split into randomized 100 image partitions to speed up data processing. Approximately 7,643 particles were picked using a 3D template picker with a 13-protofilament microtubule reference, with a separation distance between particles of 82 Å. Particles were extracted and 4 × binned (3.328 Åper pixel) then 2D classified to generate 7 dataset-derived 2D references for template picking. These dataset-derived references were used for 2D template picking to pick 51,367 particles on all 4,448 micrographs.

These particles were extracted at a box size of 720 pixels, 4 × binned to 180 pixels (3.328 Åper pixel), and protofilament sorted using an established method of supervised 3D classification (without blush regularisation) with 6 references corresponding to 11 2, 12 3, 13 3, 14 3, 15 4, and 16 4 synthetic microtubules (Cook et al., 2020; Sui and Downing, 2010). 26,851 particles corresponding to the majority, 14-protofilament, 3-start helix class were subsequently 2D classified, and class averages without proteinaceous features were discarded. The remaining 33,948 particles were subjected to 3D Refinement with helical symmetry applied (He and Scheres, 2017) using the 14 3 synthetic reference, achieving 4*×* binned Nyquist of 6.7 Å. Aligned particles were re-extracted without binning and re-refined with helical symmetry applied to 4.0 Åusing the binned map as a reference. Newly aligned particles were CTF Refined with per-particle defocus and astigmatism and subsequently polished using default parameters. CTF Refined and polished particles were re-refined with helical symmetry applied to 3.5 Åusing the prior refined map as a reference.

Aligned particles were symmetry expanded by 28× to 650,104 particles using relion particle symmetry expand and the helical parameters from the final helical refinement (twist = -25.77^*°*^, rise = 8.750 Å). A 4-dimer, 2-protofilament patch mask was generated using relion mask create with molmaps of fitted tubulin dimers in ChimeraX (Goddard et al., 2007, 2018). The patch mask was then used to perform Signal Subtraction (Bai et al., 2015) on the 28 × symmetry-expanded particles, with re-centering enabled and a new box size of 400 pixels. The signal-subtracted particles were reconstructed without angular refinement using relion reconstruct to generate a centered reference for subsequent 3D Refinements. The signal-subtracted particles were then subjected to non-helical 3D Refinement without symmetry applied, resulting in a 2.9 Åmap lacking an α- and β-tubulin register.

To register sort the particles, 4 different molmaps representing every possible lattice configuration (αβ/αβ, αβ/βα, βα/βα, βα/αβ) were made in ChimeraX at a resolution of 6 Å. The signal-subtracted particles were then sorted into these classes using an alignment-free supervised 3D Classification with a Regularisation parameter T of 200 to prioritize high-resolution information. 235,419 particles sorted into the αβ/αβ register class were then taken forward and re-refined to 2.9 Å. Particles were then subjected to per-particle CTF Refinement and rerefined and sharpened to 2.7 Å. Finally, the half-maps from the final 3D Refinement were density-modified with the Resolve CryoEM program (Terwilliger et al., 2020) from the Phenix structural biology suite (version 1.21.1-5286; Adams et al., 2010) to a resolution of 2.4 Åand resampled without smoothing in Coot by a factor of 1.3.

#### Model building and refinement

The atomic model for the 4-dimer Dm-Tb was generated by fitting separate AlphaFold2 (Jumper et al., 2021)models of *Drosophila* α-tubulin A1 (AF2 code AF-P06603-F1-model v4) and β-tubulin B1 (AF2 code AF-Q24560-F1-model v4) into the central 4 subunits (chains B,D,E,H) of the 4-dimer microtubule patch density modified map. The starting model was subject to a real-space refinement in Phenix using default parameters (Afonine et al., 2018). The refined subunit models were manually rebuilt in Coot (version 0.9.8.95; Emsley and Cowtan, 2004; Emsley et al., 2010) and were used as templates for the other copies of tubulin (chains C,I,J,K) present in the map. GDP and GTP ligands were added to corresponding inter-dimer contact sites, with coordinate magnesium ions added to all GTPs and water molecules built adjacent to the magnesium ions in chains D and H. The complete model was then subject to further Phenix real-space refinement, Coot rebuilding, and a final round of Phenix real-space refinement. All residues show strong backbone density except known disordered regions (α-tubulin residues 38-46, α-tubulin residues 441-452, and β-tubulin residues 429-449) and β-tubulin 126-129, which correspond to a surface loop *∼*20 Åfrom a lateral contact site.

#### Data visualization

All components of Fig. 3 were created in ChimeraX except for Fig. 3H, which was made in Microsoft Excel. The smooth protein chain models used in Fig. 3D&F were generated using the ChimeraX command “car style protein modeh default arrows f xsect oval width 1 thick 1”. The displacement vectors used in Fig. 3D&F were generated using pseudobond files in ChimeraX. The histograms in Fig. 3E&G were generated using the ChimeraX “crosslinks histogram” command to measure the pseudobond files. Fig. 4F&G and Fig. S5D were created in ChimeraX using the “worms” option when rendering by attribute using the RMSF values in Fig. 4H or b-factor values in Fig. S5D. Fig. S5A. Raw images and 2D classes inFig. S4 were displayed in RELION while 3D density maps were displayed in ChimeraX. Fig. S5A&E were generated in RELION, Fig. S5B was generated from the Resolve Cryo-EM job in Phenix, and Fig. S5F was generated in Coot.

#### Layer line analysis of *Drosophila* GDP-microtubule lattices

Microtubule lattice geometries of dynamic Dm-Tb microtubules were determined by taking advantage of the pseudo-helical nature of microtubules using Fourier transform-based layer line analysis. This analysis was aided by TubuleJ, a plugin created to analyze the lattice parameters of individual microtubules in Fiji (Ku et al., 2020; Schindelin et al., 2012). In this work, TubuleJ was used primarily to straighten the microtubule image using a Bessel function (Blestel et al., 2009). The determination of lattice parameters was conducted by comparison of the resulting Fourier transforms to previously published Fourier transforms of specific lattice types (Chrétien et al., 1996).

Specifically, the direction of the pf skew, θ, was determined by visual inspection of the JS and JN-S layer lines on the Fourier transform (Chrétien et al., 1996). A positive, right-handed θ was assigned if JS was closer to the equator than JN-S, and a negative, left-handed θ was assigned if JN-S was closer to the equator Fig. S6. The predicted protofilament number (N) and helix start number (S) were determined for each microtubule by qualitative comparison to known Fourier transform patterns (Chrétien et al., 1996). 13 protofilament microtubules with a 3-start helix were identified based on their Fourier transform pattern and the lack of an arrowhead pattern in the filtered microtubule image, which was generated with TubuleJ. Low resolution 3D reconstructions of individual microtubules were computed with TubuleJ to validate the predicted lattice geometries (N S) and used for illustrative purposes.

### Molecular Dynamics simulations of tubulin dimers

Models for molecular dynamics were started from structures of *Drosophila* α1/β1 and bovine α1B/β2A tubulin dimers in the GDP state. For the Bt-Tb dimer we used a structure from our previous work (Hanson et al., 2016) and for the Dm-Tb dimer we used a structure of a GDP-bound dimer (PDB ID: 7QUC) (Wagstaff et al., 2023). Missing atoms or residues were rebuilt using simple homology modeling in Visual Molecular Dynamics (VMD) (Humphrey et al., 1996). Importantly, the unstructured C-terminal tails of the tubulins were not included in the molecular dynamics simulations. Each system was solvated using TIP3P water with 10 Åpadding. Na^+^ and Cl^*−*^ were added to neutralize the charge of the system and give an ionic strength of 50 mM. NAMD 3.0 (Phillips et al., 2020) was used to perform the simulations using the CHARMM36 forcefield (Huang and MacKerell Jr, 2013). Following minimization, the systems were heated and equilibrated at a temperature of 300 K and 1 atm of pressure (NpT conditions). Simulations were performed with 2 fs timesteps by fixing hydrogens. We employed Particle Mesh Ewald (Essmann et al., 1995) for long-range electrostatics and used a 10 Åcut-off and 8.5 Åswitch distance for van der Waals interactions. After equilibration for 100 ns, we ran four independent simulations of 0.8 µs each for both Dm-Tb and Bt-Tb, resulting in 3.2 µs for each tubulin dimer.

PCA and RMSF trajectory analysis was carried out in R using the Bio3D v2.3-3 package (Skjærven et al., 2014) and visualization was again conducted using VMD.

### Allocolchicine binding assays

#### Assay set up

Samples of Dm-Tb and Bt-Tb bound to GTP and GDP were prepared from previously purified tubulin stored in BRB80 buffer. Free nucleotide was first removed from the solution of tubulin by desalting the tubulin into nucleotide-free buffer (NFB; 50 mM Hepes, 50 mM KCl, 1 mM EGTA, 1 mM MgCl_2_; pH 7.1) with a NICK desalting column (Sephadex G-50, Cytivia). The tubulin in NFB was concentrated, the concentration was measured with a concentration curve of absorbance at 280 nm, and then 1 mM of either GTP or GDP was added to the sample. This tubulin was flash frozen in small aliquots and stored at -80^*°*^C until the binding assays were conducted. Crystallized allocolchicine was kindly gifted by Tyler Heiss and Tarun Kapoor. Dilute stocks of allocolchicine were prepared by dissolving allocolchicine crystals in DMSO and the concentration of allocolchicine was determined by measuring the UV absorbance at 276 nm with an extinction coefficient of 10250 M^*−*1^cm^*−*1^ (Medrano et al., 1989).

The fluorescence emitted by binding of allocolchicine to tubulin was measured with a Cary Eclipse fluorescence spectrophotometer. The excitation wavelength used in all experiments was 310 nm and emission was collected at 425 nm. These wavelengths were chosen as they represent the peaks in multiple scans for optimal excitation (250-370 nm) and emission (350-650 nm) wavelengths for solutions of allocolchicine and tubulin in NFB. The photomultiplier tubes (PMT) voltage on the spectrophotometer was set to 650 volts and the emission signal was averaged ten times over 0.1250 seconds. An emission slit size of 10 nm was used. Experiments were conducted in opaque 384-well plates at room temperature, measured as 25^*°*^C across trials.

#### Kinetics binding assay

For the kinetics binding assay, the increase in fluorescence generated by the binding of 10 µM allocolchicine to 4 µM tubulin was measured. Fluorescent signal was collected every 20-30 seconds for the first 15 minutes of the acquisition, and then once every 15 minutes for the remaining 2 hours. The percent saturation of fluorescent intensity at each time point was calculated by dividing the intensity at time t by the maximum intensity measured in the assay, to better compare experiments preformed separately. The percent saturation data was fit to the following exponential function:

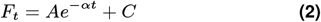

Where F_t_ is the fluorescent intensity at time t, A is the amplitude of the kinetic phase, α is the kinetic association rate, and C is the y-intercept. The y-intercept constant accounts for the slight time delay between mixing of the solution and the first measurements. Fitting was conducted using the SciPy package in Python for all of the allocolchicine assay analysis (Virtanen et al., 2020).

#### Equilibrium binding saturation assay

To estimate if the differences in the kinetic association of allocolchicine to Dm-Tb and Bt-Tb were due to differences in binding affinity, we probed the affinity of allocolchicine to the two tubulin dimer samples with an equilibrium binding saturation curve. The tubulin samples, buffers, and allocolchicine stocks used were the same materials as used in the binding kinetics assay. The fluorescence generated by the binding of varying concentrations of allocolchicine (0, 2.5, 5, 7.5, 10, 15, 25, 40 µM) to 4 µM tubulin was measured after 2 hours of equilibration to ensure full binding saturation. Five consecutive measurements taken 15 minutes apart were averaged to determine the maximum saturation for each sample on the curve, with the final measurement taken 3 hours 15 minutes after initial mixing. The data was fit to the following binding saturation curve:

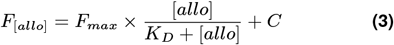

Where F_[allo]_ is the fluorescent intensity at a particular allocolchicine concentration, F_max_ is the maximum fluorescent intensity, and C is the y-intercept. The y-intercept constant in this equation accounts for the non-zero starting value resulting from non-specific fluorescent emission of tubulin in solution without allocolchicine.

### Temperature stability analysis with nano-differential scanning fluorimetry

The melting temperature and onset of unfolding temperatures of Dm-Tb and Bt-Tb were measured using a Prometheus Panta nanoDSF machine. Both Dm-Tb and Bt-Tb were diluted to 10 µM in BRB80 buffer and measured in capillaries. The temperature was steadily increased from 20^*°*^C to 85^*°*^C in 0.132^*°*^C ± 0.126^*°*^C increments. The data analysis was conducted with the Prometheus instrument software. Briefly, the ratio of absorbance at 350 nm / 330 nm was used to create the melting curve and determine the onset of unfolding temperature, while the first derivative of the curve was used to find the melting point. Three independent repeats were conducted for each tubulin type.

### Statistical testing

Statistical tests used are indicated in figure legends: equality testing and Student’s t-test. Statistical testing was conducted in Python using the SciPy statistical functions, scipy.stats, for t-tests and calculating the p-value of Z-values (Virtanen et al., 2020). A custom script was created for equality testing based on equation 4 in “Statistical Test for the Equality of Regression Coefficients” (Paternoster et al., 1998). In all tests, p-values *>* 0.05 were considered non-significant. Linear regression was carried out with weighted-least squares regression from the statsmodels modules (Seabold and Perktold, 2010).

## ACKNOWLEDGEMENTS

The authors thank Tyler Heiss and Tarun Kapoor for the synthesis of allocolchicine; Kastauv Basu and S.K. Sears at the McGill Facility for Electron Microscopy Research (FEMR), Kahn Huy Bui, Corbin Black, and Thibault Legault for support setting up cryo-EM experiments; Luke Rice for helpful discussions about the project; and Maria Genova for helpful advice about TOG-affinity chromatography. Infrastructure from the Integrated Quantitative Biology Initiative (Canadian Foundation of Innovation 33122), FEMR at McGill, the Centre de recherche en biologie structurale (CRBS), and U-M Cryo-EM was utilized in this research. The CRBS is supported by Fonds de Recherche du Québec (Health Sector) Research Centres Grant #288558. U-M Cryo-EM is grateful for support from the U-M Life Sciences Institute and the U-M Biosciences Initiative. Computational infrastructure used in this article was supported by NIH grant S10OD020011. N.V. and M.A.C. were supported by NIH grant R01GM141119. M.S. was supported by an FRQNT Doctoral Training Scholarship (https://doi.org/10.69777/317016), a Lorne-Trottier Accelerator Fellowship from McGill University, and a NSERC Canadian Graduate Scholarship – Masters. M.S. and A.N.B. were recipients of CRBS Studentship Awards from the Centre de recherche en biologie structurale, funded by Fonds de Recherche du Québec (Health Sector) Research Centres Grant #288558. A.N.B. was additionally supported by a CIHR Doctoral Fellowship. J.-F.T. holds a Canada Research Chair (Tier 2) in Structural Pharmacology (#950-229792) G.J.B. acknowledges support from Natural Sciences and Engineering Research Council of Canada (RGPIN-2020-04876), the Canadian Institutes of Health Research (PJT-148702), and McGill University.

## AUTHOR CONTRIBUTIONS

M.S. and G.J.B. conceived of the project. N.V. collected EM micrographs, processed the 3D-reconstruction, and analyzed the results for the cryo-EM experiments. A.N.B. performed mass spectrometry experiments and analyzed the data. S.C.T. wrote scripts for IRM image processing, and dynamics data fitting & analysis. W.P. performed nano-DSF experiments. D.S. performed molecular dynamics simulations and analyzed the data. M.S. performed all other experiments and analyzed the data. M.S., N.V., D.S., and A.N.B. prepared figures for data visualization. J.-F.T. contributed reagents for mass spectroscopy. M.A.C. contributed reagents for cryo-EM. G.J.B., D.S., M.A.C., and J.-F.T. secured the funding for the research. M.S., N.V., D.S., and G.J.B. wrote the manuscript with input from all authors.

## Supplementary Information

**Fig. S1.**
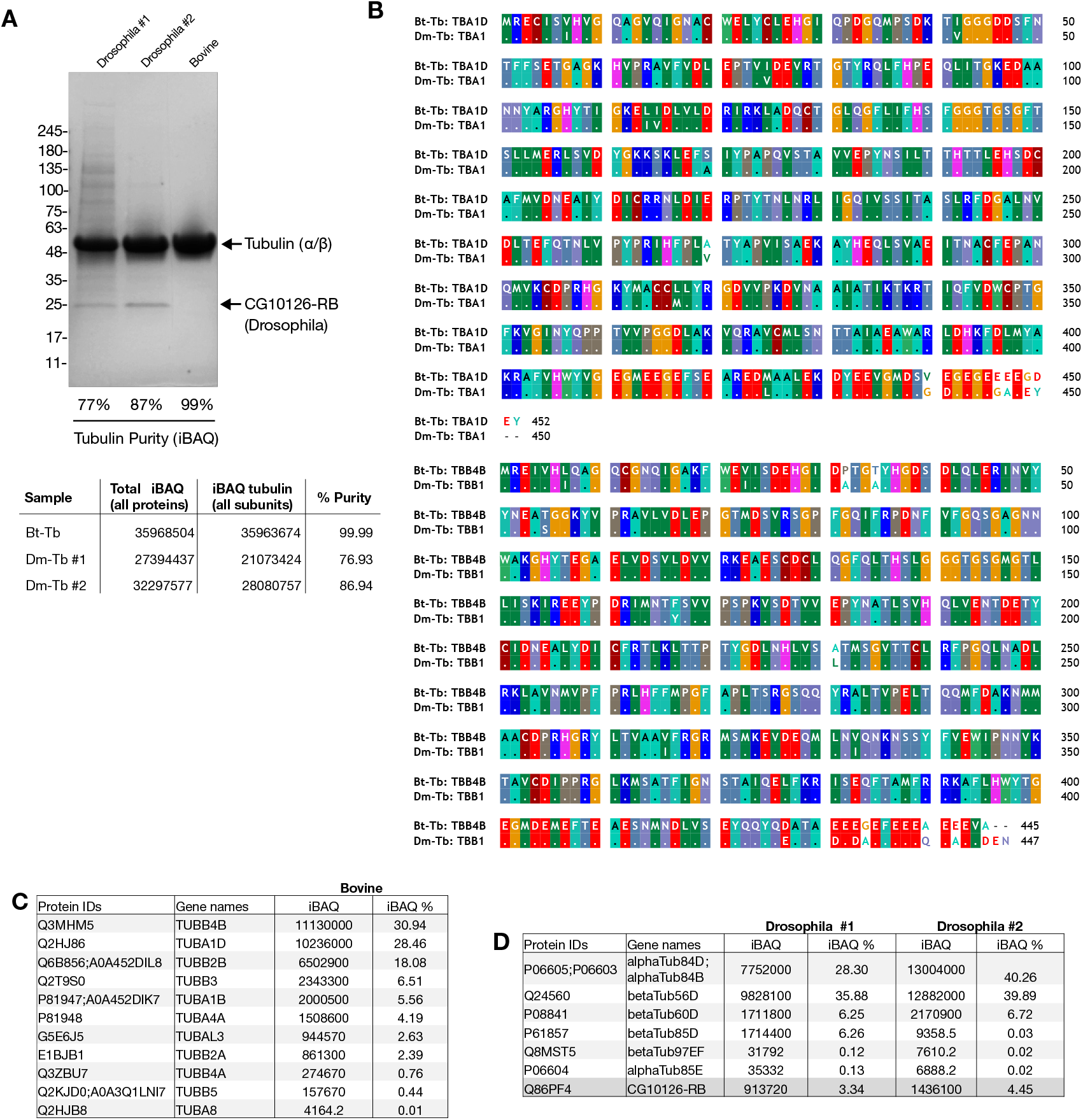
SDS-PAGE and mass spectrometry analysis of purified tubulin. Related to Fig. 1. **A)** Purified tubulin samples were resolved on SDS-PAGE and visualized with Coomassie stain. Tubulin monomers and CG10126-RB are labeled according to their migration on the SDS-PAGE gel. Tubulin purity was calculated by taking the summed iBAQ values of all tubulin proteins identified and dividing this value by the total iBAQ for all proteins identified in the mass spectrometry dataset (visualized in table below). **B)** Pairwise sequence alignment of the most abundant isotypes in the samples. For α-tubulin, the most abundant isotypes are α1 for Dm-Tb (P06603) and α1D for Bt-Tb (Q2HJ86). For β-tubulin, the most abundant isotypes are β1 for Dm-Tb (Q24560) and β4D for Bt-Tb (Q3MHM5). Sequences retrieved from UniProt and aligned in BioEdit. Conservation is indicated by colored blocks and identical residues are denoted by a period in the Dm-Tb sequence. Non-conservative substitutions are shown on a white background. **C)** Summary of mass spectrometry data for individual tubulin subunits in a representative Bt-Tb sample. **D)** Summary of mass spectrometry data for individual tubulin subunits and CG10126-RB in a representative Dm-Tb sample. (A,C-D) iBAQ values are reported directly from MaxQuant. iBAQ % represents the individual protein group’s iBAQ value divided by the summed iBAQ value for all identified proteins (excluding contaminants, only identified by site, and reverse hits).

**Fig. S2.**
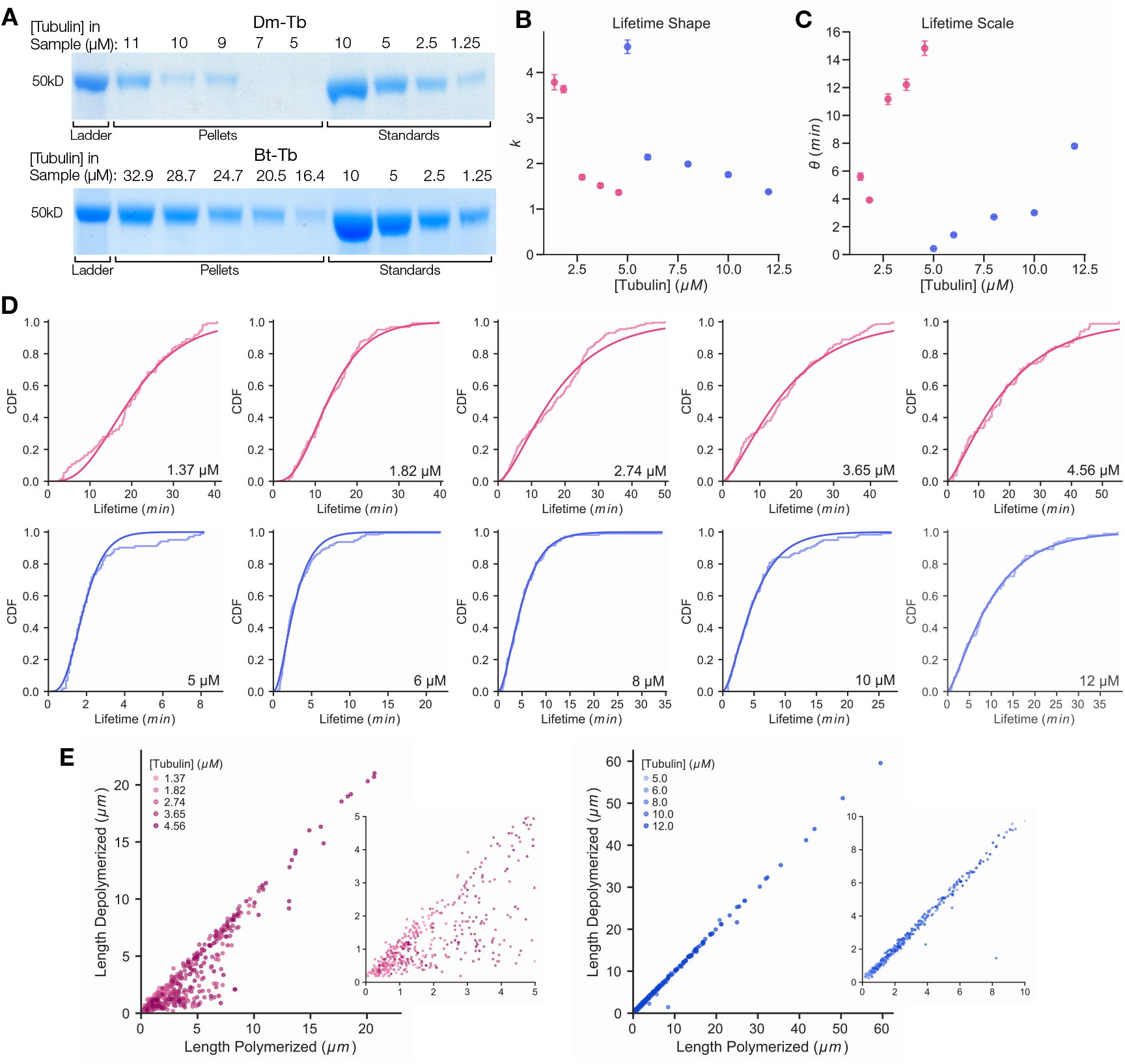
Supplementary data on lifetimes, nucleation, and rescue. Related to Fig. 2. SDS-PAGE gel of tubulin recovered in microtubule pellets for Dm-Tb (top) and Bt-Tb (bottom) after 1 hour at 36*°*C. Pellets were rarely observed for 5 and 7 µM Dm-Tb samples. Standards of known tubulin concentrations were included on every gel to control for differences in staining. **B)** Shape parameter of the Gamma distribution fits as a function of tubulin concentration for Dm-Tb (pink) and Bt-Tb (blue). **C)** Scale parameter of the Gamma distribution fits as a function of tubulin concentration for Dm-Tb (pink) and Bt-Tb (blue). **D)** Fitted Gamma functions to the cumulative distribution function (CDF) of microtubule lifetimes at different tubulin concentrations for Dm-Tb (pink) and Bt-Tb (blue). The same CDFs as shown in Fig. 2E with the means of the Gamma fits plotted in Fig. 2F. **E)** Plots of total length depolymerized before rescue or shrinkage to the seed as a function of the total length polymerized before catastrophe showing that catastrophe is randomly distributed along the shrinkage length. Data for Dm-Tb rescue events are in pink while Bt-Tb rescue events are in blue. Few Bt-Tb rescue events were observed.

**Fig. S3.**
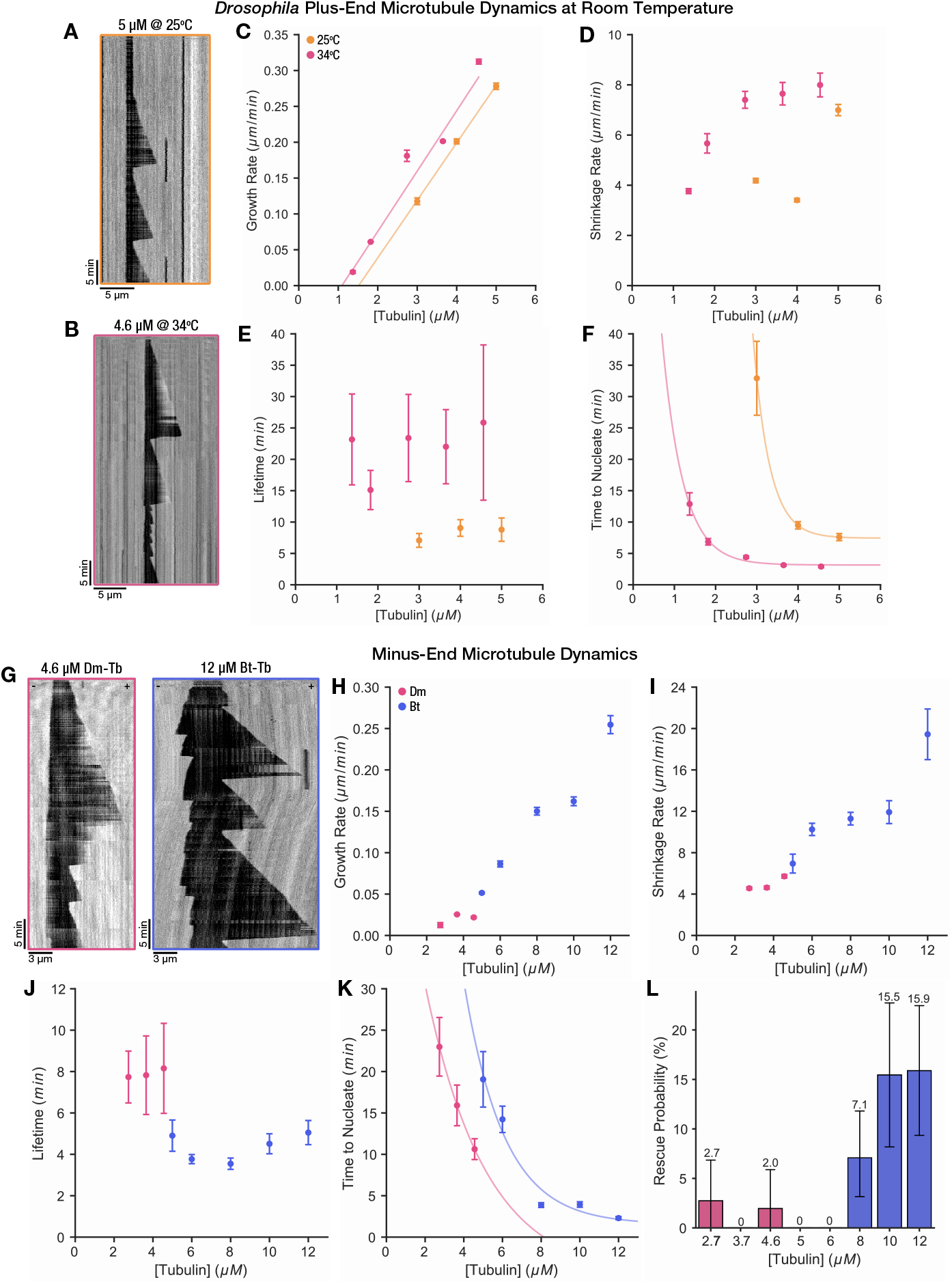
Analysis of microtubule plus-end dynamics at room temperature and minus-end dynamics at 34*°*C. Related to Fig. 1 and 2. Representative kymograph of Dm-Tb microtubule growth events at 25*°*C. **B)** Representative kymograph of Dm-Tb microtubule growth events at 34*°*C. **C-F)** Microtubule dynamics parameters for Dm-Tb at 34*°*C (pink) and 25*°*C (orange). **C)** Plot of the fitted mean and standard error of microtubule growth rate as a function of tubulin concentration. Small SE bars are not visible. **D)** Plot the fitted mean and standard error of microtubule shrinkage rate as a function of tubulin concentration. Small SE bars are not visible. **E)** Plot of the mean microtubule lifetimes as a function of tubulin concentration. Means calculated with a gamma fit and error bars represent the standard error of the fit. **F)** Plot of mean time to nucleate as a function of tubulin concentration. Means calculated with an exponential fit and error bars represent the standard error of the fit. 34*°*C data in (B-F) 34*°*C is the same as presented in Fig. 1 and Fig. 2. 25*°*C data has n = 129, 203, 177 from 3+ replicates. **G)** Representative kymographs of microtubule growth events at the minus end of Dm-Tb and Bt-Tb microtubules. No minus end growth was observed below 2.7 µM for Dm-Tb microtubules. **H)** Plot of the fitted mean and standard error of microtubule minus end growth rate as a function of tubulin concentration. Small SE bars are not visible. **I)** Plot the fitted mean and standard error of microtubule minus end shrinkage rate as a function of tubulin concentration. Small SE bars are not visible. **J)** Plot of the mean microtubule minus end lifetimes as a function of tubulin concentration. Means calculated with a gamma fit and error bars represent the standard error of the fit. **K)** Plot of mean time to nucleate at the minus end as a function of tubulin concentration. Means calculated with an exponential fit and error bars represent the standard error of the fit. **L)** Rescue probability of microtubule minus ends. Data in (G-L) are measured from the same microtubules presented in Fig. 1 and Fig. 2 (Dm-Tb: n = 73, 51, 51 from 3 replicates; Bt-Tb: n = 40, 97, 127, 110, 107 from 3 replicates).

**Fig. S4.**
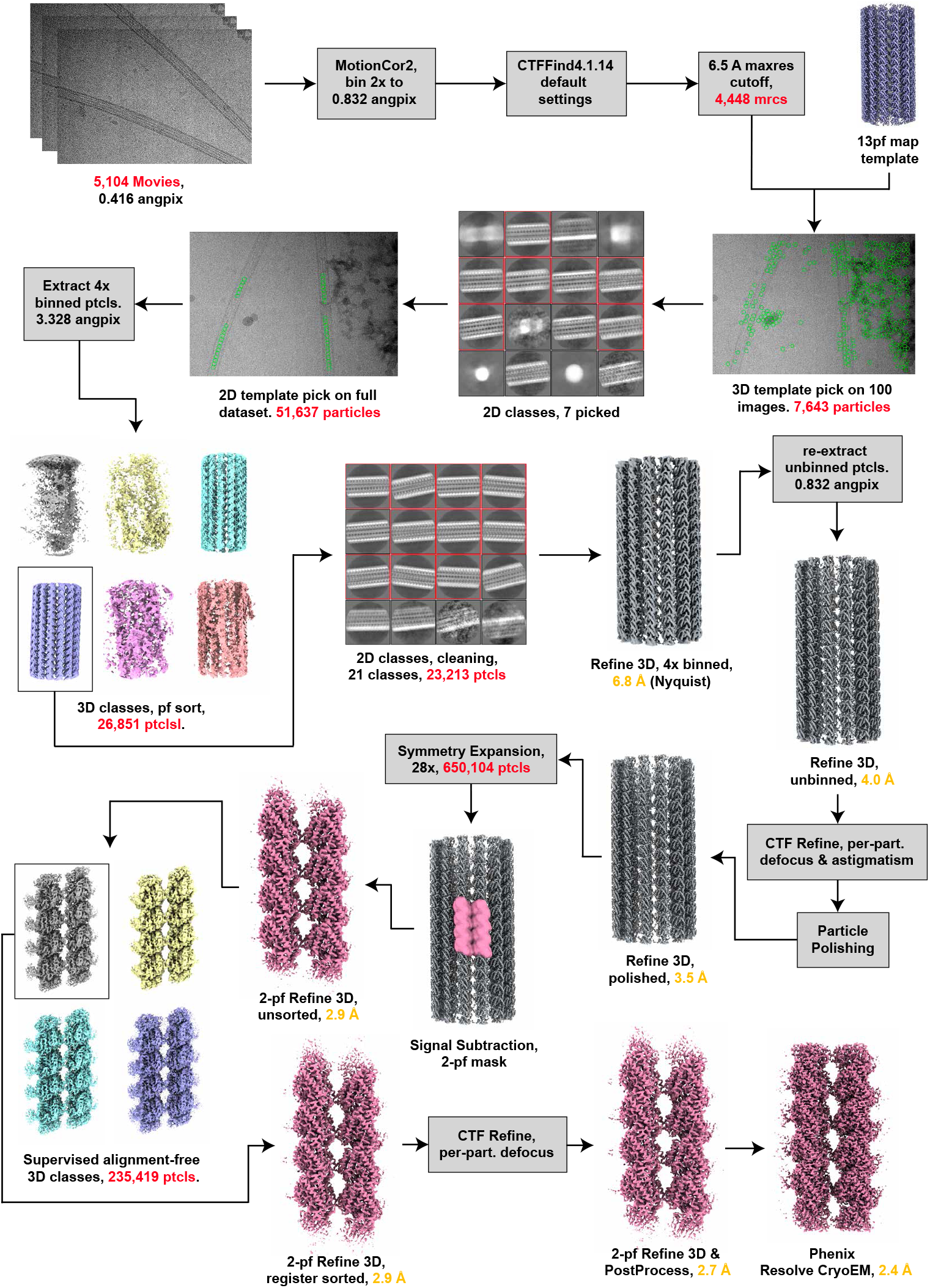
Cryo-EM data processing pipeline figure. Related to Fig. 3. Visual depiction of the workflow described in “Image processing and 3D-reconstruction” Methods sub-section. Red text corresponds to a change in image or particle number. Gold text corresponds to a change in estimated resolution. Gray boxes indicate computational steps lacking a visual output. All steps were performed in RELION-5.0-beta except for mask creation, which was partially done with molmaps made in ChimeraX, and the final density modification step, which was performed in Phenix.

**Fig. S5.**
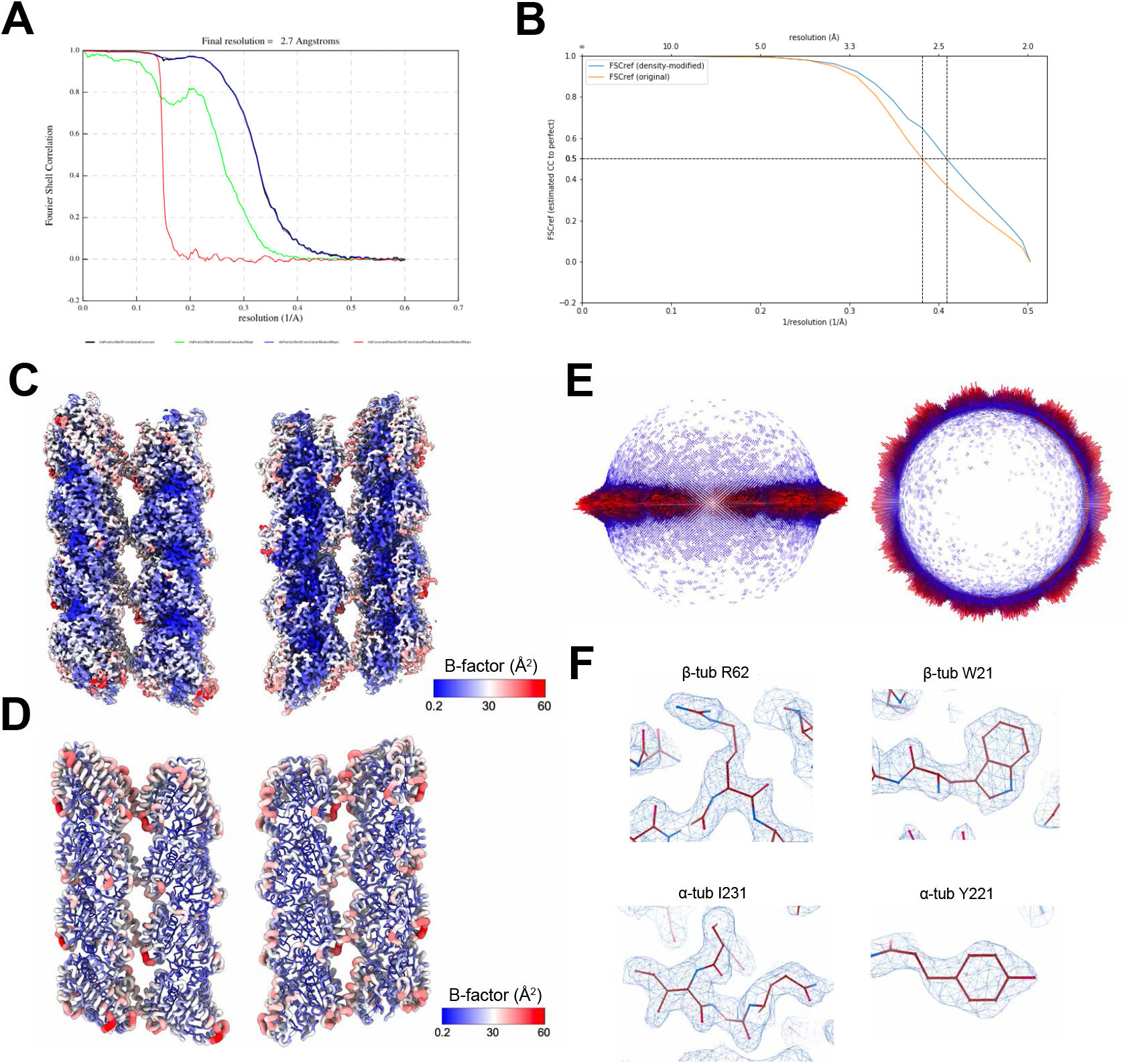
Supplementary data on the cryo-EM density map. Related to Fig. 3. **A)** FSC cross-correlation plot of the sharpened, non-modified density map generated by RELION. **B)** FSC cross-correlation plot of the density-modified map generated by Phenix. **C)** Density-modified map colored by B-factor (in Å_2_). **D)** Molecular model colored and dilated by B-factor (in Å_2_). **E)** Angular distribution of particles in final reconstruction generated by RELION. **F)** Select high-resolution side-chain densities.

**Fig. S6.**
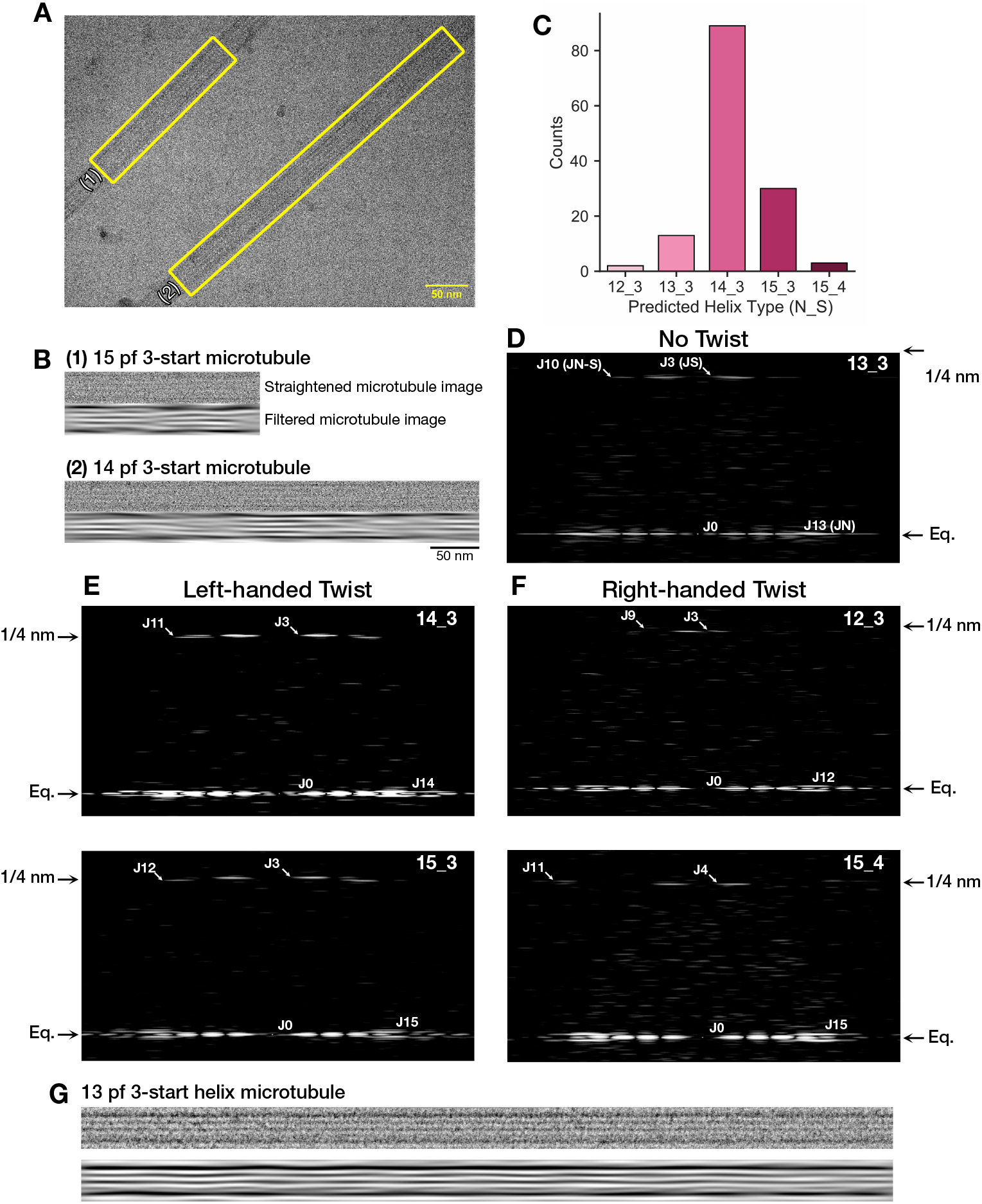
Analysis of microtubule lattice geometry by Fourier transform inspection. Related to Fig. 3. Example micrograph showing two segments picked for lattice geometry analysis. **B)** Straightened microtubule images and filtered microtubule images of the segments picked in (A). The Moiré period is measured with a plot profile on the filtered image. The 15 pf 3-start microtubule has a shorter Moiré period than the 14 pf 3-start microtubule. **C)** Predicted lattice geometry distribution for Dm-Tb microtubules analyzed by layer line analysis of microtubule Fourier transforms. Predicted helix type combines the protofilament number, N, and the helix start number, S, as both characteristics are needed to determine the geometry of the lattice. 14 3 and 15 3 lattices both feature a negative, left-handed twist. **D)** Fourier transform of 13 pf 3-start helix microtubule lattice with no pf twist. **E)** Fourier transforms of left-handed microtubule lattices: a 14 pf 3-start lattice is shown on the top and 15 pf 3-start lattice on the bottom. **F)** Fourier transforms of right-handed microtubule lattices: a 12 pf 3-start lattice is shown on the top and 15 pf 4-start lattice on the bottom. **G)** Example images of 13 pf 3-start helix microtubule straightened (top) and filtered (bottom). The horizontal lines indicate low pf skew, as 13 pf microtubules are known for their lack of twist.

**Fig. S7.**
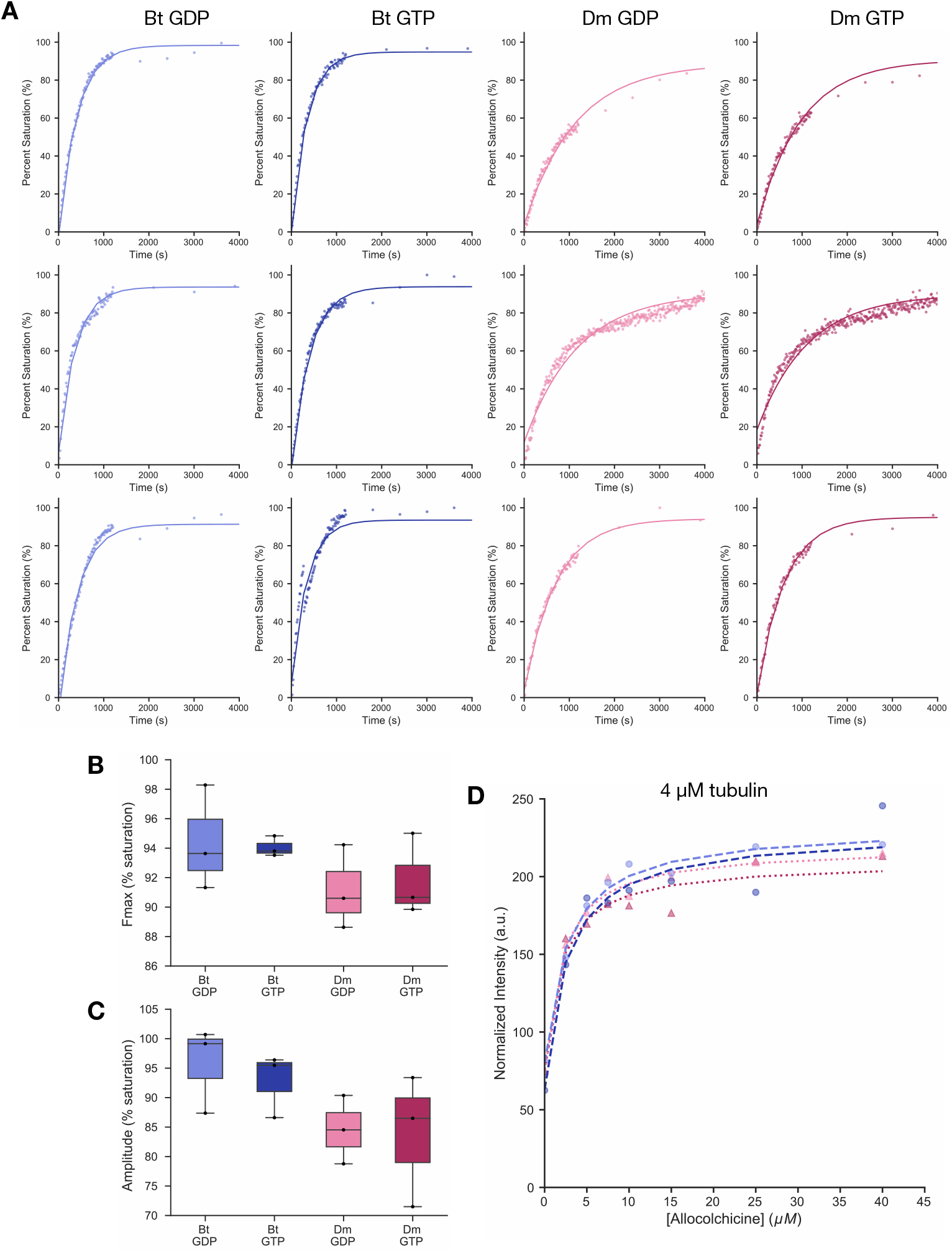
Allocolchicine binding assays raw data and fits. Related to Fig. 5. A) Raw data and fitted curves for each individual measurement of allocolchicine binding kinetics to 4 µM each Bt-Tb and Dm-Tb. **B)** Maximum fluorescence (Fmax) from the kinetic binding curve of each individual experiment in (A). **C)** Amplitude of the kinetic binding curve of each individual experiment in (A). **D)** Binding saturation curve used to determine the dissociation constant of varying concentrations of allocolchicine from 4 µM Bt-Tb and Dm-Tb bound to GTP and GDP (N = 1). The fitting of curves in (A) and (D) are described in the methods.

**Table 1.** Cryo-EM data collection, refinement, and validation table. Related to Fig. 3.

| Data collection and processing |  |
| --- | --- |
| Microscope | Titan Krios G4i |
| Voltage (kV) | 300 |
| Detector | K3 Summit |
| Magnification | 105,000 |
| Electron exposure ($e^-/\text{\AA}^2$) | 50.5 |
| Exposure rate ($e^-/\text{pixel/s}$) | 12 |
| Calibrated pixel size; superres/binning ($\text{\AA}$) | 0.416/0.832 |
| Defocus range ($\mu\text{m}$) | 0.5-2.0 |
| Symmetry imposed | C1 |
| Movies (no.) | 5,104 |
| Initial particle images (no.) | 51,637 |
| Final particle images (no.) | 235,419 |
| Map resolution ($\text{\AA}$) | 2.7 |
| FSC threshold | 0.143 |
| Density-modified map resolution ($\text{\AA}$) | 2.4 |
| Density-modified FSC threshold | 0.5 |
| Map sharpening factor ($\text{\AA}^2$) | -35 |
| Map resolution range ($\text{\AA}$) | 2.3-3.3 |
| Helical symmetry (whole 14-pf MT map) |  |
| Single subunit rise ($\text{\AA}$) | 8.75 |
| Helical twist ($^\circ$) | -25.77 |
| Refinement |  |
| Initial models (AlphaFold ID) |  |
| α-tubulin | AF-P06603-F1-model_v4 |
| β-tubulin | AF-Q24560-F1-model_v4 |
| Model resolution ($\text{\AA}$) | 2.4 |
| FSC threshold | 0.5 |
| Model composition |  |
| Non-hydrogen atoms | 27,212 |
| Protein residues | 3,436 |
| Waters | 8 |
| Ligands | 4 $\text{Mg}^{2+}$ |
| Nucleotides | 4 GTP, 4 GDP |
| B factors ($\text{\AA}^2$) | |
| Proteins | 14.71 |
| Waters | 4.01 |
| Ligands | 3.05 |
| Nucleotides | 9.21 |
| R.M.S. deviations |  |
| Bond lengths ($\text{\AA}$) | 0.003 |
| Bond angles ($^\circ$) | 0.607 |
| Validation |  |
| MolProbity score | 1.89 |
| Clashscore | 8.3 |
| Poor rotamers (%) | 2.78 |
| Ramachandran plot |  |
| Favored (%) | 97.51 |
| Allowed (%) | 2.49 |
| Disallowed (%) | 0 |
| Rama-Z score | 0.97 |
| EMRinger score | 3.81 |

